# An empirical comparison of univariate versus multivariate methods for the analysis of brain-behavior mapping

**DOI:** 10.1101/2020.04.13.039958

**Authors:** Maria V. Ivanova, Timothy J. Herron, Nina F. Dronkers, Juliana V. Baldo

## Abstract

Lesion symptom mapping (LSM) tools are used on brain injury data to identify the neural structures critical for a given behavior or symptom. Univariate lesion-symptom mapping (ULSM) methods provide statistical comparisons of behavioral test scores in patients with and without a lesion on a voxel by voxel basis. More recently, multivariate lesion-symptom mapping (MLSM) methods have been developed that consider the effects of all lesioned voxels in one model simultaneously. However, very little work has been done to empirically compare the advantages and disadvantages of these two different methods. In the current study, we provide a needed systematic comparison of 5 ULSM and 8 MLSM methods, using both synthetic and real data to identify the potential strengths and weaknesses of both approaches. We tested power and spatial precision of each LSM method for both single and dual (network type) anatomical target simulations across anatomical target location, sample size, noise level, and lesion smoothing. Additionally, we performed false positive simulations to identify the characteristics associated with each method’s spurious findings. Simulations showed no clear superiority of either ULSM or MLSM methods overall, but rather highlighted specific advantages of different methods. No single method produced a thresholded LSM map that exclusively delineated brain regions associated with the target behavior. Thus, different LSM methods are indicated, depending on the particular study design, specific hypotheses, and sample size. Overall, we recommend the use of both ULSM and MLSM methods in tandem to enhance confidence in the results: Brain foci identified as significant across both types of methods are unlikely to be spurious and can be confidently reported as robust results.

## 1 INTRODUCTION

Throughout the 19th and much of the 20^th^ century, systematic clinical observations of neurologic patients along with post-mortem autopsy remained the main method for establishing brain correlates of cognitive functioning (Damasio & Damasio, 1989; Dronkers, Ivanova, & Baldo, 2017; Luria, 1980). The advent of modern neuroimaging methods in the 1970s greatly enhanced the ability to determine neural foundations of cognition, as the actual lesion site could be identified in-vivo with unprecedented, continuously improving precision. In the early 2000s, an increase in computing power along with new statistical procedures brought lesion-symptom mapping (LSM) to a new, more advanced level. Instead of relying on single-case studies or viewing regions of lesion overlap in patients with a common syndrome, analysis of large group studies with continuous behavioral data became possible. Specifically, the mass-univariate LSM (ULSM) method, such as the original voxel-based lesion symptom mapping (VLSM; Bates et al., 2003), provides statistical comparisons of behavioral test scores across patients with and without a lesion on a voxel by voxel basis. Voxels that show significant differences for a particular behavior or symptom are inferred to be critical for the behavior under examination. ULSM methods complement functional neuroimaging studies in healthy participants, by testing the necessity of particular brain areas for a particular behavior, thereby demonstrating the crucial causal link in brain-behavior relationships (Bates et al., 2003; Karnath, Sperber, & Rorden, 2018; Rorden, Karnath, & Bonilha, 2007; Vaidya, Pujara, Petrides, Murray, & Fellows, 2019). Contemporary ULSM methods provide a fundamental shift in broadening our understanding of brain-behavior relationships, both confirming (Baldo, Arevalo, Patterson, & Dronkers, 2013) and challenging previously held beliefs about key neural structures for different cognitive functions (Baldo et al., 2018; Dronkers, Wilkins, Valin, Redfern, & Jaeger, 2004; Ivanova et al., 2018; Mirman et al., 2015).

More recently, new multivariate lesion-symptom mapping (MLSM) methods have been developed as an alternative to ULSM. The principle difference between ULSM and MLSM methods is that MLSM considers the entirety of all lesion patterns in one model simultaneously. This is in contrast to the parallel, independent analysis of lesion patterns on a voxel by voxel basis performed with ULSM models. While some papers have argued that MLSM methods should be superior to ULSM methods (DeMarco & Turkeltaub, 2018; Mah, Husain, Rees, & Nachev, 2014; Pustina, Avants, Faseyitan, Medaglia, & Coslett, 2018; Zhang, Kimberg, Coslett, Schwartz, & Wang, 2014), many of these arguments have been presented theoretically without rigorously comparing the two approaches (see also Sperber, Wiesen, & Karnath, 2019). Below, we review issues that impact the validity of brain-behavior inferences made with both ULSM and MLSM methods and set the stage for our current comprehensive evaluation of different LSM techniques.

One issue that affects both ULSM and MLSM methods is that lesion distributions in stroke (the most frequently studied etiology with LSM techniques) are influenced by the vascular anatomy and are thus non-randomly distributed in the brain (Mah et al., 2014; Phan, Donnan, Wright, & Reutens, 2005; Sperber & Karnath, 2016; Xu, Jha, & Nachev, 2018). The non-random distribution of lesions limits analysis of certain brain areas that are rarely affected in stroke (e.g., the temporal pole). An associated issue that potentially affects both ULSM and MLSM methods is that neighboring voxels have a higher probability of being lesioned together, as strokes never affect just one voxel. The ULSM approach is potentially susceptible to this spatial autocorrelation, because it assumes independence of lesioned voxels throughout the brain, as thousands of independent tests (non-parametric, t-tests, or linear regressions) are carried out serially in the affected voxels. While independence of tests is not an assumption of MLSM methods per se (since only one multivariate model incorporating all the lesion patterns is tested), lack of sufficient spatial distinction is an issue. In other words, if two voxels are always either damaged together or always spared, it is not possible to differentiate their unique contribution to the observed deficits with any LSM method. Another related concern that affects both ULSM and MLSM methods is differential statistical power across voxels/regions of the brain. For example, a voxel in which 50% of patients have a lesion has more power than a voxels where only 10% of patients have a lesion (Pustina, Avants, Faseyitan, Medaglia, & Coslett, 2018). Cumulative effects of non-random lesion distribution, autocorrelation across voxels, and differential power distribution can potentially lead to bias in spatial localization of critical regions, as significant clusters are “diverted” towards the most frequently damaged regions (Inoue, Madhyastha, Rudrauf, Mehta, & Grabowski, 2014; Xu et al., 2018) and potentially along vascular patterns (Mah et al., 2014; Sperber et al., 2019). The open question is, to what degree do these biases occur with different ULSM and MLSM methods, and if there are biases, how are they ameliorated by sample size, method choices and output interpretation?

One critique of ULSM by Mah and colleagues (2014) suggested that ULSM analyses mis-localized foci by an average of 16 mm, but their model did not include lesion volume as a covariate in their analysis. The importance of using lesion volume as a nuisance covariate in LSM has been a standard recommendation for several years (Baldo, Wilson, & Dronkers, 2012; DeMarco & Turkeltaub, 2018; Haan & Karnath, 2018; Price, Hope, & Seghier, 2017; Sperber & Karnath, 2017). In addition, Mah et al. used a minimum lesion load per voxel of < 1% in their ULSM analyses, which is far below the standard recommendation of 5 – 10% (Baldo et al., 2012). Moreover, the displacement maps using synthetic data in Mah et al. showed single voxels (i.e. a single voxel leading to a specific deficit), which is an oversimplified and exclusively theoretical case that does not occur naturally. Furthermore, when damage to an anatomical region was used as a synthetic behavioral score in their study, the score was binarized rather than continuous, likely further reducing spatial resolution. Finally, Mah et al. did not provide the spatial bias values for MLSM, so it was not possible to directly compare the results to ULSM.

In another simulation study critiquing accuracy of ULSM (Inoue et al., 2014), lesion volume was included as a covariate, but the authors again used binarized synthetic behavioral scores (a deficit was indicated when 20% of voxels in the target parcel were damaged) and did not apply a minimal lesion load threshold. Also, the results in this study were predominantly analyzed with False Discovery Rate (FDR) based thresholding. This method of correction for multiple comparisons has been discontinued for some time in the ULSM literature, as it frequently leads to an increase in false positives (Baldo et al., 2012; Kimberg, Coslett, & Schwartz, 2007; Mirman et al., 2018). Also, accuracy of mapping was not systematically explored across sample sizes. Finally, the lesion data for this study came from highly heterogenous etiologies (stroke, traumatic brain injury, encephalitis), contrary to standard recommendations for any LSM study (Haan & Karnath, 2018).

Most prominently, Sperber and Karnath (2017) empirically demonstrated that ensuring a sufficiently large minimal lesion load threshold as well as including lesion volume as a covariate have additive effects on reducing spatial bias. They have also shown that adding “anatomical bias” vectors to the model (calculated from inter-voxel relationships in the anatomical data) markedly reduced spatial bias up to 7.0 mm. In their study, spatial bias was calculated for single voxels, with displacement of larger clusters expected to be smaller. Furthermore, the lesion volume correction to enhance accuracy of localization has been strongly recommended for at least some MLSM approaches (DeMarco & Turkeltaub, 2018), again highlighting that MLSM methods are not immune to these types of spatial biases. In another simulation study, Sperber and colleagues (2019) showed that a common support vector regression-based MLSM method was also susceptible to mislocalization along the brain’s vasculature, even after applying a correction for lesion volume, and that this displacement error was actually higher than that observed for a ULSM method. However, since displacement was determined for single voxels in a single axial slice, these spatial biases require further exploration to fully understand their impact on LSM results with real behavioral data.

The most comprehensive simulation study to date by Pustina et al. (2018) showed that even one of the most advanced MLSM algorithms, sparse canonical correlation analysis for neuroimaging (SCCAN), exhibited spatial bias in the results. Here, a superiority of SCCAN using synthetic data was consistently demonstrated, but only when compared to the univariate analyses with FDR-based thresholding. In the same paper, the ULSM results obtained with more appropriate thresholding using permutation-based and Bonferonni Family-Wise Error Rate (FWER) corrections, were comparable to SCCAN results across a number of spatial indices (Pustina et al., 2018). Moreover, Pustina et al. (2018) did not include lesion size as a covariate in the ULSM analysis, running counter to standard recommendations for ULSM and potentially biasing the comparison (Baldo et al., 2012; Sperber & Karnath, 2017). Furthermore, limited spatial metrics were used as measures of accurate mapping in comparing LSM methods, and most of these metrics produced similar levels of performance for all methods tested. For example, while the dice index (measure of overlap between two regions) was shown to be significantly higher for SCCAN compared to a non-parametric Brunner-Munzel version of ULSM, values were very low in nearly all cases with every method (predominantly < 0.5 and often < 0.2) rendering the statistical advantage uninformative. Also, results of statistical comparisons across different sample sizes for other spatial metrics were not provided (see Figure 4, p.161, Pustina et al., 2018). In short, the degree to which spatial bias affects ULSM versus MLSM methods has not yet been systematically and rigorously tested across a wide range of realistic LSM choices and under a wide range of metrics of spatial accuracy.

Another known issue for in LSM is the ability of different LSM techniques to detect complex relationships and dependencies in the data (i.e., networks). Some papers have argued that MLSM should be better than ULSM at detecting multivariate relationships between lesion location and behavioral deficits, as MSLM takes into account all the voxels simultaneously in a single model (DeMarco & Turkeltaub, 2018; Mah et al., 2014; Pustina et al., 2018; Zhang et al., 2014). However, the superior ability of MLSM methods to identify networks remains to be proven. This is in part due to the strong regularization (e.g., sparse vs. dense solutions) and additional assumptions (e.g., restriction on possible locations of solutions) required in order to solve a single, massively under-determined, multivariate system of equations (typically with 1 patient per 100 or 1000 voxels). Mah et al. (2014) claimed that MLSM resulted in higher sensitivity and specificity compared to ULSM in detecting a two-parcel fragile network (when the synthetic score was based on the maximal lesion load among a set of anatomical regions). However, as described above, synthetic behavioral scores were binarized for their analysis, the statistical threshold used was not specified, and there was no quantification of the differences in spatial bias between ULSM and MLSM. Pustina et al. (2018) showed an advantage of MLSM over ULSM in detecting an extended network (‘AND’ rule; when the synthetic score was based on the average lesion load among a set of anatomical regions) consisting of 3 parcels. However, there was no significant advantage of the MLSM over ULSM in detecting other types of 2- and 3-parcel networks when a proper FWER correction was included. Previous ULSM studies with real data have repeatedly shown that ULSM methods are able to detect spatially distinct regions in a network (Akinina et al., 2019; Baldo et al., 2018; Gajardo-Vidal et al., 2018; Mirman et al., 2015), but it is still unknown how much power ULSM has vs. MLSM in detecting multifocal behavioral determinants.

A different issue that is restricted to ULSM methods is the problem of multiple comparisons. Since ULSM requires multiple statistical comparisons to be carried out simultaneously, and because the statistical tests as noted above are not independent, this raises the possibility of false positive errors. Modern ULSM methods use a conservative, permutation-based FWER correction, a nonparametric resampling approach to significance testing, which sets the overall probability rate of false positives across all of the results, while making almost no assumptions about the underlying data distributions (Hayasaka & Nichols, 2003; Nichols & Holmes, 2001). Permutation-based FWER provides the most stringent and robust form of correction for multiple comparisons, providing an optimal balance between false positives and false negatives (see Kimberg et al., 2007; Mirman et al., 2018). Thus, the problem of multiple comparisons in ULSM is addressed with the use of appropriate statistical corrections. Interestingly, a recent study also suggested that correction for multiple comparisons might be required for at least some MLSM methods (Sperber et al., 2019).

There are also a number of statistical challenges that are specific to MLSM. First, since all measurement units are considered in one model simultaneously, it is unclear how to estimate the necessary sample size to have sufficient power other than by intensive simulations. Second, existing MLSM models require (hyper)parameter selection (e.g., cost, gamma, sparseness), but procedures for selecting those parameters are not fully transparent to most clinical researchers or are likely to be computationally too expensive to be generated automatically (e.g., using nested cross validation; DeMarco & Turkeltaub, 2018; Pustina et al., 2018). It is not clear to what extent changing these parameters alters the obtained pattern of results, and why different parameter values are sometimes chosen for analyses within the same paper (e.g., see Fig. 7, p. 164, Pustina et al., 2018). The issue of transparency in how hyperparameters control solution regularization is a serious one, as these parameters determine the *a priori* biases of the type of solutions that any given MLSM method uses. Third, to an average user, it is not clear how most models arrive at the outcome (i.e., it is not as straightforward as the general linear model approach in ULSM). This lack of familiarity will impact the interpretation of results and limit the ability of the user to know when the method should not be used. Last, models used in MLSM can be computationally costly, which limits the ability to do post-hoc computations such as random subsampling in order to test the stability of a given solution.

To summarize, there are a number of theoretical concerns for both ULSM and MLSM methods. Some of these concerns raised originally with respect to ULSM (e.g., spatial bias and autocorrelation, differential statistical power) are actually concerns for both ULSM and MLSM methods and require further elucidation with respect to both approaches. Moreover, efficient controls already exist for both ULSM and MLSM methods that can be implemented to minimize the biasing effect of lesion physiology (e.g., lesion size correction, minimum lesion overlap threshold, and most recently, “anatomical bias” vectors). The theoretical concerns about ULSM methods being less able to detect networks of brain regions (as opposed to a single target region) have not been experimentally confirmed. Other issues such as parameterization details are specific to MSLM methods. Properties of new LSM models require further delineation and comparative evaluation in order to assess the mapping power and accuracy under varying conditions. To date, the comparisons of ULSM and MLSM in the literature have been limited and when they are contrasted, a sub-standard version of ULSM is often implemented without proper correction, leading to an unfavorable impression of ULSM (Inoue et al., 2014; Pustina et al., 2018; Zhang et al., 2014). Finally, neither ULSM nor MLSM methods have been properly explored with respect to the incidence of false positive results.

The current paper aimed to address these gaps in the LSM literature and provide a comprehensive appraisal of several versions of ULSM and MLSM methods with a large stroke lesion dataset, using both synthetic and real behavioral data, across a range of relevant parameters. Synthetic data were used to test the spatial accuracy of 13 different LSM methods. Obtained results were compared across different anatomical target locations, sample sizes, noise levels, lesion mask smoothing values, types of networks, and false positive simulations. We used a number of different distance- and overlap-based spatial metrics as indices of mapping accuracy. We also compared performance of the 13 LSM methods using real behavioral data (language scores) with multiple demographic and sampling covariates, along with subsampling to check the stability and agreement across methods. Our goal was to provide the first comprehensive comparison of ULSM and MLSM methods, in order to afford guidance on selecting the most appropriate LSM method(s) for a particular study with a specific lesion dataset.

## 2 METHODS

### 2.1 Participants

For the simulation analyses, lesion masks from 340 chronic left hemisphere stroke patients were obtained from two different sources: our Northern California stroke dataset (n = 209, NorCal) and the Moss Rehabilitation stroke dataset provided with the open-source LESYMAP software (n = 131, LESYMAP, Pustina et al., 2018). Synthetic behavioral scores were based on lesion load to different cortical areas (described further below).

For the analysis of real behavioral data, we analyzed language data and lesion masks from a subset of patients in the NorCal database (n = 168; 36 female) who completed behavioral testing and met the following inclusion criteria: History of a single left hemisphere stroke (including both embolic and hemorrhagic etiologies), pre-morbidly right-handed (based on the Edinburgh Handedness Inventory), native English speaker (English by age 5), minimum high school or equivalent education (i.e., 12 years), in the chronic stage of recovery (at least 12 months post-stroke) at the time of behavioral testing, no other neurologic or severe psychiatric history (e.g., Parkinson’s, dementia, schizophrenia), and no substance abuse history. The mean age of this subset of patients was 61.0 years (range 31-86, SD = 11.2), mean education was 14.9 (range 12-22, SD = 2.4), and mean months post-stroke was 51.4 (range 12-271, SD = 54.0). All patients were administered the Western Aphasia Battery (WAB, Kertesz, 1982, 2007), which classified 47 patients with anomic aphasia, 45 with Broca’s aphasia, 6 with conduction aphasia, 4 with global aphasia, 1 with transcortical motor aphasia, 3 with transcortical sensory aphasia, 14 with Wernicke’s aphasia, and 48 patients who scored within normal limits (i.e., overall WAB language score of ≥ 93.8 points out of 100). This latter group included patients with very mild aphasic symptoms, such as mild word-finding difficulty.

### 2.2 Behavioral data

Data for the LSM analyses with real behavioral scores were derived from 168 patients in the NorCal stroke dataset. Patients were tested on the WAB (Kertesz, 1982, 2007), which consists of several subtests measuring a wide range of speech and language functions. Here, we analyzed the most reliable and least-confounded speech-language scores on the WAB, which index three distinct language domains: speech fluency, single-word auditory comprehension, and verbal repetition. All patients signed consent forms and were tested in accordance with the Helsinki Declaration.

### 2.3 Imaging and lesion reconstructions

Real lesion masks were obtained from two different sources as detailed previously. All lesion masks were reconstructed from MRI or CT data acquired during the chronic phase of stroke (at least 2 months post-stroke). Detailed information about data acquisition, lesion reconstruction, and normalization procedures for the NorCal and LESYMAP datasets can be found in Baldo et al. (2013) and Pustina et al. (2018), respectively. The lesion masks were converted to standard MNI space with a 2mm isovoxel resolution. The overlay of patients’ lesions from the two different databases is shown in Figure 1. Mean lesion volume was 119.6cc for the NorCal dataset (range 0.1-455, SD = 97.9) and 100.0cc for the Moss Rehab dataset (range 5.2-371.4 SD=82.2).

**Figure 1.**
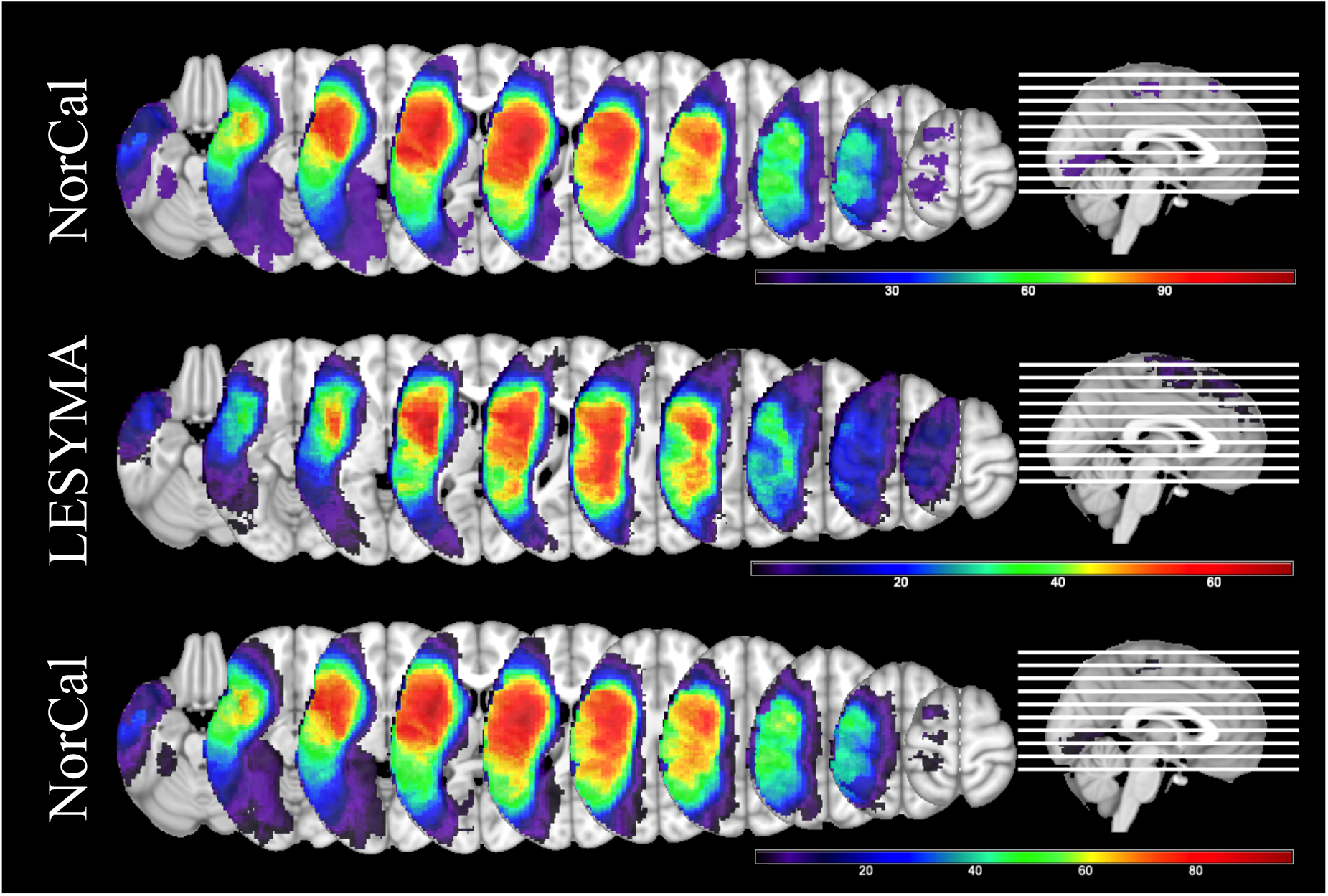
Lesion overlays for the three datasets. Top – NorCal (*n* = 209, coverage range 5 – 117). Middle – LESYMAP (*n* = 131, coverage range 5 – 68). Bottom – subset of NorCal used in the analysis of real behavioral data (*n* = 168, coverage range 5 – 96).

### 2.4 LSM methods

In the current study, we compared 5 ULSM and 8 MLSM methods, using both synthetic and real behavioral data. All ULSM and MLSM methods discussed in this paper were implemented in a downloadable MATLAB script (freely available at https://www.nitrc.org/projects/clsm/) that was based on the original VLSM software developed by Stephen Wilson (Bates et al., 2003; Wilson et al., 2010).

#### 2.4.1 ULSM methods

All ULSM variants were conducted using a linear regression with a voxel lesion value as the dependent variable (lesioned or not), the continuous behavioral data as the independent variable, and lesion volume as a covariate (Wilson et al., 2011; Baldo et al., in press). Linear regression was chosen because it is flexible, popular and applicable to multiple data types (categorical or continuous) with multiple covariates. Also, the synthetic behavioral data contained linear effects, similar to other recent simulation studies (e.g., Pustina et al., 2018). Both the synthetic and real behavioral datasets were well controlled with respect to outliers; all the z-scores were within 3 SD of the mean.

To correct for multiple comparisons, we used permutation-based thresholding, which is a non-parametric approach to FWER correction that randomly permutes behavioral scores and which records the relevant t-value or maximal cluster size for each permutation with the final threshold set at *p* = .05 (Kimberg et al., 2007; Mirman et al., 2018). Unlike some previous methodological LSM papers, we did not evaluate the performance of the ULSM method with FDR-correction, given that it is a fundamentally inappropriate correction for lesion data (Baldo et al., 2012, in press; Mirman et al., 2018). Finally, the use of permutation-based significance testing with all of the ULSM methods protects against inflated false positives that can accompany non-Gaussian noise that may appear in the real data (all synthetic behavioral noise was Gaussian).

ULSM results were generated with five different non-parametric FWER thresholding approaches that are commonly used in contemporary ULSM studies:

1. Maximum statistical t-value (ULSM T-max). This statistic corresponds to the most standard and conservative version of the permutation-based FWER-correction;
2. 125^th^-largest t-value (ULSM T-nu=125). This statistic corresponds to the 125^th^ largest voxelwise test statistic (n = 125 corresponds to 1 cm^3^ when working with 2 mm-sided voxels; see Mirman et al., 2018);
3. Cluster-size thresholding with a fixed voxel-wise threshold of *p* < .01 (ULSM T-0.01);
4. Cluster-size thresholding with a fixed voxel-wise threshold of *p* < .001 (ULSM T-0.001);
5. Cluster-size thresholding with a fixed voxel-wise threshold of *p* < .0001 (ULSM T-0.0001).

#### 2.4.2 MLSM methods

##### MLSM

methods included 2 monolithic regression methods and 6 data reduction methods. For all MLSM methods, lesion volume was regressed out of both the behavioral and the lesion variables (DeMarco & Turkeltaub, 2018). We normalized both lesion and behavioral data to standard deviation of 1 and centered the behavioral data (to mean 0), as is customary for multivariate methods to optimize regression estimation. Finally, as with ULSM methods, permutation testing was also used to threshold and identify significant voxels (*p* < 0.05, using the maximum statistical value obtained).

##### Support Vector Regression (SVR)

differs from ordinary multivariate regression in a number of ways, as it computes a solution based on many voxels’ lesion status given the relatively small number of patients. First, it incorporates two regularization hyperparameters that help the regression model keep the model’s parameter values small (to avoid overfitting), while at the same time controlling the model’s prediction accuracy by partially ignoring small fitting errors. Second, SVR incorporates a radial basis kernel function that implicitly projects the lesion data into a high dimensional space in order to help model fitting succeed, in part by allowing some nonlinear effects to be incorporated into the model. SVR has been used in several previous LSM studies (Ghaleh et al., 2017; Griffis, Nenert, Allendorfer, & Szaflarski, 2017; Zhang et al., 2014). We used the SVR routine encoded as part of the SVR-LSM package (https://github.com/dmirman/SVR-LSM; DeMarco & Turkeltaub, 2018) that uses fixed hyperparameters previously tuned to work well in LSM (Zhang et al., 2014).

The second monolithic MLSM regression method was **Partial Least Squares (PLS)**, which jointly extracts dual behavior and lesion factors that maximize the variance between behavior and lesion locations in a single step. PLS algorithms (and closely related canonical correlation algorithms) have been developed extensively in bioinformatics for use in genetics where there is a similar “wide” data structure: there are far more genes/voxels to be considered in a regression solution than there are subjects providing such data (Boulesteix & Strimmer, 2006). PLS has also previously been used in LSM (Phan et al, 2010). However, we used a basic version of PLS regression based on the singular value decomposition function (Abdi, 2010; Krishnan, Williams, Randal, & Abdi, 2011) because it is known to be both a fast and reliable regression technique over wide data sets containing highly correlated variables, which is important given the number of simulations (using permutation testing) that were run. Although this version of PLS regression is known to produce “dense” solutions (Mehmood, Liland, Snipen, & Sæbø, 2012), resulting in (overly) large clusters, another important consideration for including it here as an exemplar of this class of algorithms is that it is easy to generalize basic PLS regression to integrate multiple target behaviors simultaneously (Abdi, 2010); an inviting prospect for investigating behavioral test batteries used to assess patient populations.

##### Data reduction methods

Three different types of MLSM data reduction methods were tested in the current study: Singular Value Decomposition, Logistic Principal Component Analysis, and Independent Component Analysis (described below). These data reduction methods reduce the spatial dimensionality of the lesion data first without considering behavior. Lesion status of thousands of voxels is reduced to a number of spatial lesion components that is fewer than the number of patients in the analysis. This results in a more tractable system of equations to solve, and the components’ estimated weights are then transformed back into spatial maps, relating brain areas to the behavior being investigated. In other words, lesion status in all voxels for each patient are replaced with sums of weighted lesion components for that patient.

##### Singular Value Decomposition (SVD)

(“svd”, MatLab v.7) (Ramsey et al., 2017) identifies an ordered set of orthogonal spatial components, each a weighted mixture of all voxels inside the lesion mask, that explain as much lesion variance across all voxels with as few of the ordered components as possible. These components can be linearly combined to reconstruct each patient’s lesion mask. Our preliminary trials found that most lesions were well-reconstructed when 90% of the cumulative variance was accounted for. Thus, in the data-reduction version of MLSM we used the number of components (approximately equal to half the number of patients) required to explain 90% variance.

The second data reduction method we used was a **Logistic Principal Component Analysis (LPCA)** (Schein, Saul, & Ungar, 2003), which iteratively identifies a set of ordered spatial components whose lesion incidence maps are orthogonal under the logistic function (Siegel et al., 2016). Given the better fit of method-to-lesion data types, trials with lesion masks showed that approximately the first 40 LPCA components were able to reconstruct patient lesion masks quite well (typically dice > 0.99). Thus, the number of components we used for the LPCA data reduction was the number of patients capped at 40 components.

The third data reduction method was an **Independent Component Analysis (ICA)** (FastICA v2.5 as used by Hyvärinen & Oja, 2000) which is a generalization of PCA. ICA in this context estimates independent linear mixtures of lesion incidence voxel data that are the most non-Gaussian sources found within the data. We used the default cubic function as the fixed-point non-linearity for finding components under coarse iterations first and then used a hyperbolic tangent function (default setting) for fine iterations in order to reflect the bounded nature of lesion data. For ICA, we used the same number of components as with LPCA (maximum of 40), in order to see if ICA can outperform LPCA given its usefulness in other spatial dimension reduction applications in neuroimaging (Calhoun, Liu & Adali, 2009).

For the three data reduction approaches described above, an elastic net linear regression (“glmnet” package; Qian, Hastie, Friedman, Tibshirani, & Simon, 2013; Tibshirani et al., 2010) was performed with the target behavior (real or simulated) as the dependent variable along with the data reduced spatial lesion components and the lesion size covariate. We used two different elastic net regressions to see if either is superior in producing accurate or reliable LSM maps: one near to a pure lasso (which we will call “L1”) case (95% L1 penalty mixed with a 5% L2 penalty) and one with the opposite mixture (95% L2 & 5% L1), a ridge (“L2”) case. The elastic net regressions use cross-validation to solve for the penalty hyperparameter that best fits the data. Thus, there were 6 data reduction approaches in total: SVD-L1, SVD-L2, LPCA-L1, LPCA-L2, ICA-L1, and ICA-L2.

### 2.5 Simulations with synthetic behavioral data

Three sets of simulations were performed with all ULSM and MLSM methods using synthetic behavioral scores and real lesion masks: single anatomic target, dual anatomic targets, and zero anatomic target (i.e., false positive simulation). In each simulation, we varied several factors (described below for each simulation) in a fully crossed manner in order to systematically compare effect sizes and significance across the different ULSM and MLSM methods. For each simulation analysis, the specified number of lesion masks were randomly selected from one of the two datasets (i.e. not mixing NorCal and LESYMAP masks together). For all analyses, we only included voxels in which at least 5 patients had lesions, and which had statistical power ≥ .1 at *p* < .01 (Hsieh, Block & Larsen, 1998). We also performed behavioral value outlier scrubbing at |*z*|≥ 3.0, which was particularly important at higher behavioral noise levels.

Synthetic behavioral scores (also called “artificial” or “fake” in the literature) for the single and dual anatomical target simulations were derived from the lesion load to the target anatomical parcels (or ROIs). For simple single target simulations, the synthetic behavioral score was calculated as the fraction of the target anatomical parcel that was spared (see next section for more details), i.e., the synthetic score was directly proportional to the lesion load of that anatomical parcel. Use of synthetic behavioral scores allows one to determine how well the different LSM methods are able to localize behavior, since we know the ground truth – exactly to which region in the brain it should localize to (i.e., the target anatomical parcel) (Pustina et al., 2018). To create target anatomical parcels, we used ROI masks of grey matter areas in the left middle cerebral artery region from FSL’s version of the Harvard-Oxford (H-O) atlas, thresholded at 50% incidence. We used 16 such parcels that had 5% or greater lesioned area within at least 25% of the lesion masks. To create a set of smaller parcels, each of these 16 parcels was divided into two sections along the axis of maximal spatial extent. Two of the subdivided parcels failed to intersect the lesion mask sufficiently according to the above criteria, rendering a total of 30 smaller parcels (see Figure 2). We specifically chose larger ROIs (similar to Mirman et al., 2018), because small ROIs are unlikely to be accurately identified in patients who generally have much larger lesions than focal fMRI activation areas. Further, even assuming that fMRI properly delineates the size and location of specific functional areas, in the chronic stage of stroke recovery, these functional areas are likely to be altered by neural reorganization (Kiran, Meier, & Johnson, 2019; Stefaniak, Halai, & Ralph, 2019).

**Figure 2.**
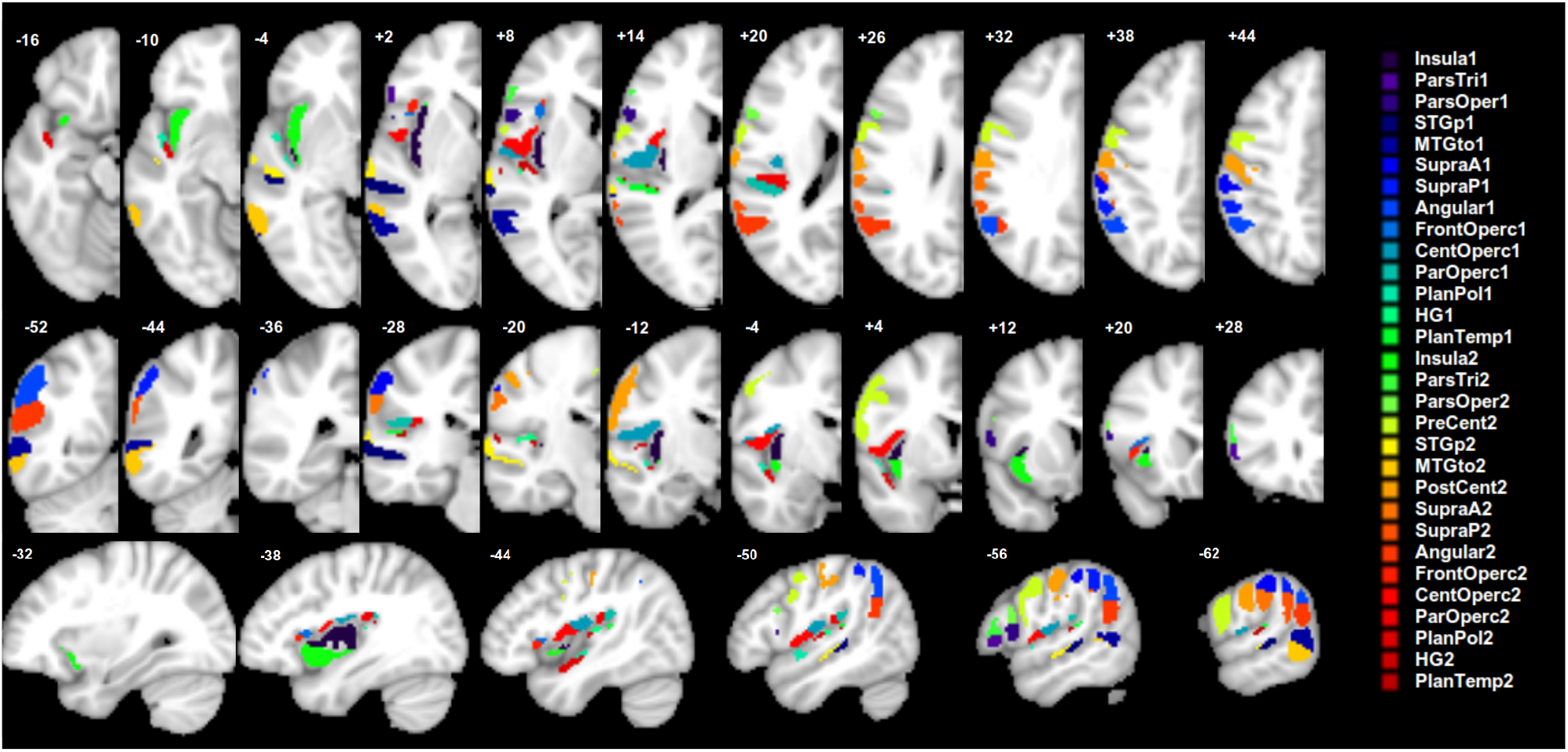
Target anatomical parcels (n = 30) used to generate synthetic behavioral scores.

#### 2.5.1 Single anatomical target simulations

In the single anatomical target simulations, the synthetic behavioral data for each patient were calculated as 1 minus the lesion load (fraction of the target anatomical parcel covered by the patient’s lesion), before noise was added. Accordingly, a score of 1 indicated “perfect performance” – complete sparing of the target parcel by the lesion, and a score of 0 indicated “complete impairment” – full coverage of the target parcel by the lesion. For the simulations, we varied the following factors in a fully crossed manner (obtaining all possible combinations of these factors):

- Number of patients (n=32, 48, 64, 80, 96, 112, 128). Additional simulations were run with n = 144-208 patients from the NorCal dataset for descriptive purposes only. Patients were randomly selected from one dataset on each run.
- Behavioral noise level (0.00, 0.36, or 0.71 SD of normalized behavioral scores). Behavioral noise level was a fixed additive level of Gaussian noise at the specified fractional level of the mean across all target measure’s standard deviations.
- Lesion mask smoothing (0 mm or 4 mm Gaussian FWHM). The smoothed mask values fell between 0 and 1, and the total mask weight was kept constant. All LSM methods could handle continuous values via regression.
- Size of parcels (16 larger or 30 smaller as described above).
- Anatomical target parcel (see Figure 2 for list).

#### 2.5.2 Dual anatomical target simulations

In the dual anatomical target simulations, two spatially distinct target parcels were used to simulate a minimal brain “network.” We considered three different types of dual-target networks:

- *Redundant network* in which the behavioral score is reduced only if both target parcels are lesioned (one minus the minimum lesion load of the two parcels is used to generate the synthetic behavioral score);
- *Fragile network* in which the behavioral score is reduced if either target is lesioned (one minus the maximal lesion load of the two parcels is used to generate the synthetic behavioral score; corresponds to what Pustina et al., 2018 called the “OR” rule for generating multi-region simulations and approximates the partial injury problem described in Rorden, Fridriksson, & Karnath, 2009);
- *Extended network* in which the behavioral score is reduced proportionally to the overall damage to the two regions, which is similar to the single target simulation except the parcel is divided into two spatially separate components (one minus the average lesion load of the two parcels is used to generate the synthetic behavioral score; corresponds to what Pustina et al., 2018 call the “AND” rule).

For the dual-target simulations, we only analyzed results with the larger 16 parcels, moderate behavioral noise level (0.36 SD), and lesion smoothing at 4 mm FWHM. We tested all 120 pairwise combinations of the target parcels for each type of network. The number of patients was varied systematically from n = 64 to 208. We did not use n = 32 or n = 48, because preliminary results showed a lack of power with this sample size. We only used the NorCal dataset for this analysis for consistency across these simulations.

#### 2.5.3 False positive simulations

In the false positive simulation, the behavioral variable consisted of pure Gaussian white noise. Any clusters detected in this simulation are thus false positives. This simulation was run 35,000 times per cohort size (from n = 32 to 128). There was no distance or power to detect, but rather the spatial characteristics of the false positive clusters were analyzed, including the size, number of clusters, and maximum statistical value for ULSM (MLSM values are rather arbitrary given data normalization). We also evaluated which LSM methods produced false positive clusters in a correlated fashion in order to see how much independence the methods have from each other. The number of patients included in the false positive simulation along with lesion mask smoothing values (0mm vs 4mm Gaussian FWHM) were systematically varied.

#### 2.5.4 Measures of LSM success in simulation analyses

As a measure of statistical power in both single and dual-target simulations, we calculated the proportion of trials that yielded any significant (above-threshold) LSM statistical values. For the false positive simulations, we calculated the proportion of trials that yielded above threshold LSM statistic and the number and the size of the false positive clusters.

To gauge the accuracy of an LSM method in our single and dual anatomical target simulations, we calculated two types of accuracy measures: distance and overlap. For each measure, the target anatomical parcel, whose lesion load was used to generate the specific synthetic behavioral score, was compared to the LSM output map (LSM thresholded statistical map). If an LSM method fully identified the underlying substrate, then the target parcel and the LSM output map should overlap perfectly. Our measures of accuracy were selected to provide a comprehensive evaluation of the precision of the different LSM methods.

Distance-based measures were used to compare a single target anatomical parcel to the LSM output. Three different indices of the *LSM output map* position were used: mean centroid location (COM_LSM_), mean centroid location weighted by statistical values (wCOM_LSM_) and maximum statistic location (Max_LSM_). Likewise, two different indices of the *target anatomical parcel* position were employed: mean centroid location (center of mass; COM_target_), and nearest location to the LSM output map position (Closest_target_). This resulted in 6 possible measures used to evaluate the accuracy of mapping of single target location (i.e, distance between target parcel and LSM output). See Figure 3 for an illustration of these different indices. Distance-based measures were not used for evaluation of dual-target simulations, because distances between the target parcel and LSM output could not be calculated unambiguously in this instance.

**Figure 3.**
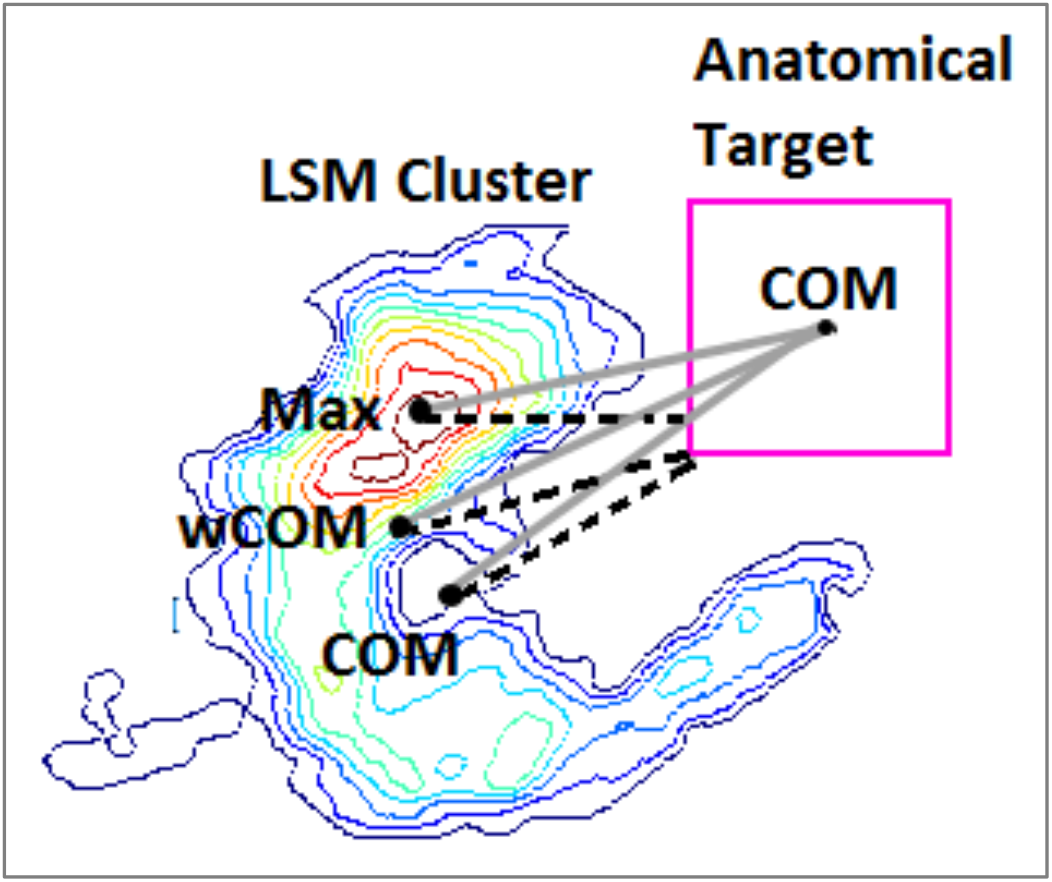
Distance-based measures of accuracy. The six distances computed in the single target simulations are represented by distinct lines: solid grey for COM_target_ distances, and dashed black for Closest_target_ distances. The LSM map is represented as a contour heat map, and the hypothetical anatomical target is a pink square.

Overlap-based measures included overlap and weighted overlap metrics between target anatomical parcel(s) and LSM output map. First, we used the *dice coefficient*, which is a simple measure of overlap between two binary maps, with 1 being perfect overlap between the two maps and with zero voxels falling outside of the overlapping area, and 0 being no overlap between the two. While the dice coefficient remains a standard means of measuring mapping accuracy, there are two clear limitations to this measure. First, the dice coefficient is greatly dependent on the size of the parcels being compared, with larger parcels generating a larger dice index compared to smaller regions, even when the relative overlap is smaller (Pustina et al., 2016). Second, the actual statistical values of the LSM output are not taken into account in the calculation of the dice index. To account for this latter limitation, we also computed a *one-sided Kuiper (OSK) distribution difference* (Rubin, 1969) between statistically significant LSM values inside the target anatomical parcel(s) versus outside. This measure compares the LSM statistical values outside the target (C) to those inside the target (B) plus it assigns zero values to target areas not reaching threshold in the LSM output (A) (see Figure 4). The idea is that we want to reward finding LSM hotspots inside the anatomical target(s) while ignoring lower values in LSM clusters that are outside the target. OSK values range from −1 to 1 with 1 representing that the entire target is covered with LSM statistical values higher than all those outside the target; −1 representing that the lowest LSM values (or none at all) are inside the target, and 0 representing no difference in LSM statistic distributions inside vs. outside the target anatomical parcel.

**Figure 4.**
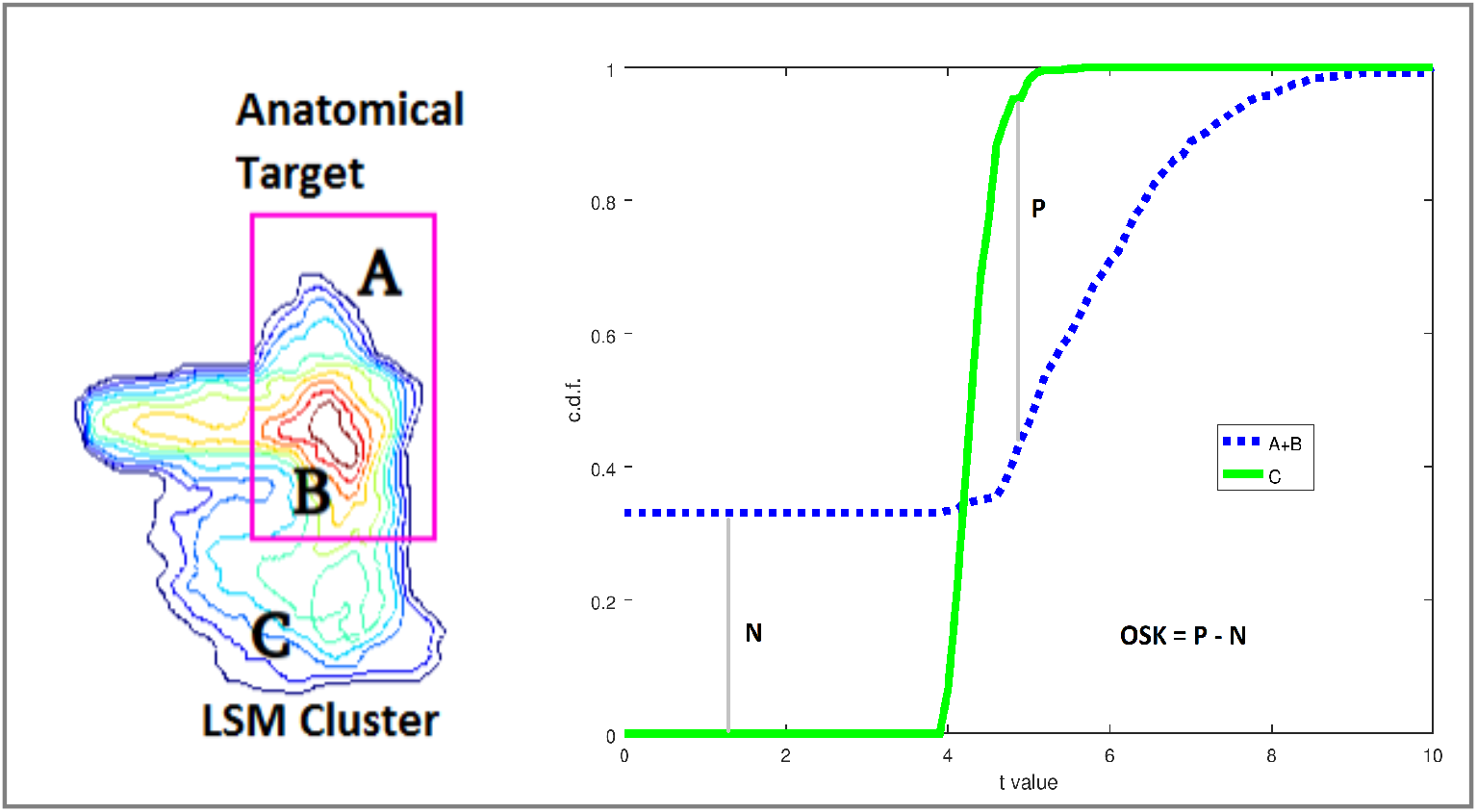
One-sided Kuiper (OSK) measure of weighted overlap. Left: The LSM statistic values inside the target (B) are aggregated with 0’s from areas inside the target not intersecting the LSM cluster (A) and both are compared with LSM values outside the target (C). Right: the cumulative distribution function (CDF) from LSM values Inside (A+B) vs. Outside (C) the target are compared and scored by subtracting the maximal CDF differences (P for C > A+B and N for C < A+B) from each other. To get a sense of how OSK values represent distributional separation, two one-dimensional standard Gaussian distributions being separated by *d* units obey the approximating equation OSK = 0.3778 • *d* – 0.0092 • *d*^3^, for *d* from [-4 to 4].

### 2.6 LSM analysis with real behavioral scores

In addition to the simulation analysis with synthetic behavioral scores, we also compared ULSM and MLSM methods using real behavioral scores. These data were generated from a subset of patients in the NorCal dataset (n = 168) who met specific inclusion criteria (described above in the participants section) and who were tested on speech fluency, single-word auditory comprehension, and repetition subtests from the WAB (Kertesz, 1982, 2007). In these analyses, in addition to lesion volume, we also covaried for age, education, gender, months post-stroke (log-transformed), test site (referring to one of the two locations where behavioral data were collected), image resolution (referring to the resolution of the original scan: low or high), and overall aphasia severity (WAB AQ minus the target subscore). The last covariate allowed us to account for overall aphasia severity, while focusing on the specific language function. Lesion smoothing was done at 4 mm FWHM. The minimum power per voxel was 0.25 at *p* < .001 for a *d*’ of 1.

We also performed 100 repetitions of the same 3 analyses but with 75% subsampling (n = 126) of the original full cohort in order to determine the stability of the solutions for each given LSM method. We considered the original LSM map generated with the full cohort (n = 168) to be the “target parcel” in this case and correspondingly analyzed the stability of the results obtained on subsequent runs (n = 126) with the same metrics we used to analyze the single target simulation results. We then tested whether there were significant differences between the different LSM methods with respect to the stability of the results produced.

### 2.7 Statistical analyses

Simulations with synthetic behavioral data were performed in a fully crossed manner: 6072 runs for the single target simulation and 10800 runs for the dual-target simulation, generating 13 LSM maps for each run. From each run’s LSM map we collected 4 datatypes: power (a binary indicator for the presence of *any* above threshold statistical values no matter the location); overlap (dice scores of LSM map with anatomical target(s)); statistic-weighted overlap (OSK, Figure 4), and, for the single target case, six distance measures (Figure 3) between the LSM map cluster(s) and the anatomical target.

Analysis of important differences between each simulation’s factors was established using mixed between/within ANOVA applied to each fully-crossed dataset type where we treated anatomical target parcel location as the random factor. We used high-speed ANOVA software (CLEAVE, nitrc.og.projects/cleave) to compute partial omega squared (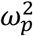) effect sizes (Olejnik & Algina, 2004) for factors and first-order interactions. Typically, weak, moderate, and strong partial omega effect sizes are taken to be 0.2, 0.5, and 0.8, respectively (Keppel, 1991). We used partial omega squared cutoff of 0.35 to restrict ourselves to reporting moderate or strong effects (unless otherwise noted). Finally, we note that the distance, dice and OSK values often did not conform to a Gaussian distribution as required by ANOVA. In these instances, we used a Kumaraswamy distribution (Jones, 2009) to model and then transform datasets into an approximately symmetric unimodal form (α=β=2) prior to ANOVA. In the presentation of results, we focus specifically on the effect of different LSM methods on the outcomes and their interactions with other factors. Additionally, we also looked at the different distance-based spatial accuracy metrics as factors and explored whether they yielded similar accuracy estimates or not. In the results we concentrate specifically on effect sizes rather than significance testing, given that large scale simulations produce so much data, that even tiny differences can be significant (Kirk, 2007; Schmidt & Hunter, 1995; Stang, Poole, & Kuss, 2010).

The same ANOVA analyses were performed on the subsample analysis of the real behavioral data by using the full analysis LSM maps as targets for their respective LSM variant (e.g., SVR target for SVR LSM analysis). The only addition was that we included the maximum statistic location of the target ROI as another centroid to anchor 3 more distances per LSM map, because these target ROIs (the n = 168 maps) already have statistics.

## 3 RESULTS

### 3.1 Single anatomical target simulations

#### 3.1.1 Statistical Power

For single anatomical target simulations, we first evaluated the statistical power of the 13 different LSM methods to detect significant voxels (see Table 1 for mean simulation values across all factor levels). Overall, all LSM methods demonstrated power > 80% with a sample size of 48 or larger, but power varied substantially across LSM Methods (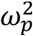 = .71). ULSM methods demonstrated the greatest power to detect significant voxels. MLSM methods required more participants to achieve comparable power (Figure 5, top panel). Sample Size (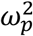 = .79) and Behavioral Noise Level (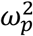 = .74) strongly impacted power across all LSM methods: larger sample size and less behavioral noise led to a higher probability of detecting significant voxels. Also, first order interactions between these three terms demonstrated a large effect size (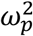 ranged from .61 to .73). At higher behavioral noise levels, a larger sample size was required to achieve similar levels of power (Figure 5, middle panel). Power was disproportionately reduced for MLSM methods relative to ULSM at higher levels of behavioral noise, with the exception of SVR (Figure 5, bottom panel).

**Figure 5.**
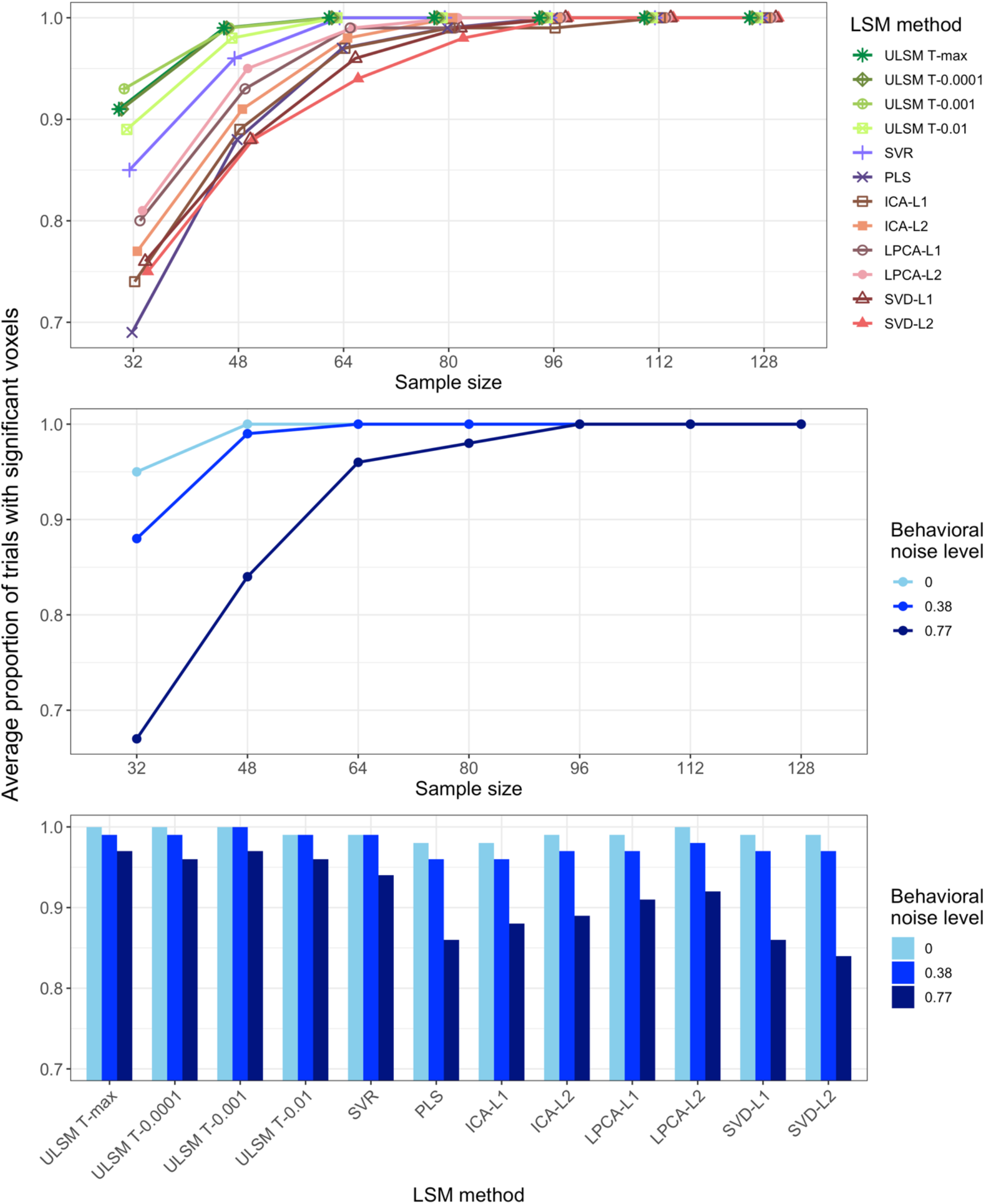
Power (average proportion of trials detecting significant voxels) as a function of the LSM method and sample size (top panel), behavioral noise level and sample size (middle panel), behavioral noise level and LSM method (bottom panel).

**Table 1.** Mean simulation values for all manipulated factors for single anatomical target simulations: percentage of trials where significant voxels were detected (Power (%)), average displacement error across all distance-based metrics (Displacement (mm)), Dice index, and One-sided Kuiper distribution statistic (OSK statistic: −1 worst to +1 best).

| LSM Method | ULSM<br>T-max | ULSM T-nu=125 | ULSM<br>T-0.0001 | ULSM T-0.001 | ULSM<br>T-0.01 | SVR | PLS | ICA-L1 | ICA-L2 | LPCA-L1 | LPCA-L2 | SVD-L1 | SVD-L2 |
| --- | --- | --- | --- | --- | --- | --- | --- | --- | --- | --- | --- | --- | --- |
| Power (%) | 98.4 | -* | 98.5 | 98.8 | 98.1 | 97.2 | 93.2 | 93.9 | 95.0 | 95.8 | 96.4 | 93.9 | 93.3 |
| Displacement (mm) | 6.1 | 6.7 | 6.4 | 6.9 | 7.6 | 6.0 | 12.9 | 12.3 | 13.1 | 12.3 | 11.4 | 8.3 | 8.0 |
| Dice index | 0.12 | 0.08 | 0.09 | 0.07 | 0.05 | 0.14 | 0.03 | 0.05 | 0.04 | 0.03 | 0.03 | 0.04 | 0.05 |
| OSK statistic | 0.43 | 0.56 | 0.52 | 0.60 | 0.68 | 0.29 | 0.54 | 0.20 | 0.35 | 0.59 | 0.61 | 0.65 | 0.66 |

| Sample size | 32 | 48 | 64 | 80 | 96 | 112 | 128 | 208** |
| --- | --- | --- | --- | --- | --- | --- | --- | --- |
| Power (%) | 82.9 | 94 | 98.4 | 99.4 | 99.8 | 99.9 | 99.9 | 100 |
| Displacement (mm) | 10.2 | 9.4 | 9.0 | 8.9 | 8.8 | 8.7 | 8.7 | 9.8 |
| Dice index | 0.10 | 0.08 | 0.07 | 0.06 | 0.05 | 0.05 | 0.05 | 0.03 |
| OSK statistic | -0.10 | 0.36 | 0.50 | 0.57 | 0.63 | 0.67 | 0.71 | 0.74 |

| Behavioral Noise Level | 0 | 0.38 | 0.77 |
| --- | --- | --- | --- |
| Power (%) | 99.2 | 97.9 | 91.9 |
| Displacement (mm) | 8.9 | 9.1 | 9.2 |
| Dice index | 0.05 | 0.06 | 0.08 |
| OSK statistic | 0.62 | 0.55 | 0.37 |

| Mask Smoothness | 0 mm | 4 mm |
| --- | --- | --- |
| Power (%) | 96.3 | 96.4 |
| Displacement (mm) | 9.2 | 8.9 |
| Dice index | 0.06 | 0.05 |
| OSK statistic | 0.49 | 0.54 |

| Number of Parcels | 16 | 30 |
| --- | --- | --- |
| Power (%) | 96.4 | 96.3 |
| Displacement (mm) | 9.4 | 8.9 |
| Dice index | 0.08 | 0.05 |
| OSK statistic | 0.38 | 0.59 |

| Anatomical Parcels | Mean | SD | Max | Min | Quart<br>1 | Quart<br>3 |
| --- | --- | --- | --- | --- | --- | --- |
| Power (%) | 96.3 | 2.7 | 99.6 | 88 | 95.3 | 98.3 |
| Displacement (mm) | 9.1 | 2.6 | 15.0 | 5.0 | 6.3 | 10.7 |
| Dice index | 0.06 | 0.04 | 0.14 | 0.01 | 0.03 | 0.09 |
| OSK statistic | 0.51 | 0.27 | 0.82 | -0.37 | 0.38 | 0.70 |

Note. \* - Power not calculated due to a technical error.
\*\* - Indices provided for informational purposes only and not included in the statistical analyses.

#### 3.1.2 Spatial accuracy using distance-based metrics

The spatial accuracy of the different LSM methods using distance-based metrics was evaluated (see Table 1 for mean displacement error values). There was a main effect of LSM Method on accuracy (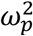 = .84). ULSM methods demonstrated numerically higher mean accuracy across all output map locations and target locations, ranging from 6-7.5 mm error. Average accuracy across MLSM methods (excepting SVR) ranged from 8-13 mm. Across sample sizes, the conservative ULSM methods (T-max and T-nu=125) and SVR produced the most accurate maps. Figure 6 shows the spatial displacement error based on mean Max_LSM_ to Closest_target_ (these two metrics in combination provide the highest accuracy estimates; more on this below) of the different LSM methods at different sample sizes. As can be seen in Figure 6, spatial displacement error varied as a function of sample size across all LSM methods, but the interaction between Sample Size and LSM Method did not reach the preselected effect size. Overall, larger Sample Size (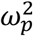 = .63) and Mask Smoothing (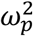 = .36) had independent positive effects on the accuracy of all LSM methods. Additionally, accuracy of LSM methods varied across the different anatomical parcels: The Rolandic operculum was detected most accurately, and the post central region was detected least accurately (see Figure 7 for displacement error across different anatomical parcels and LSM methods and Figure 8 for LSM output maps for two regions across different LSM methods).

**Figure 6.**
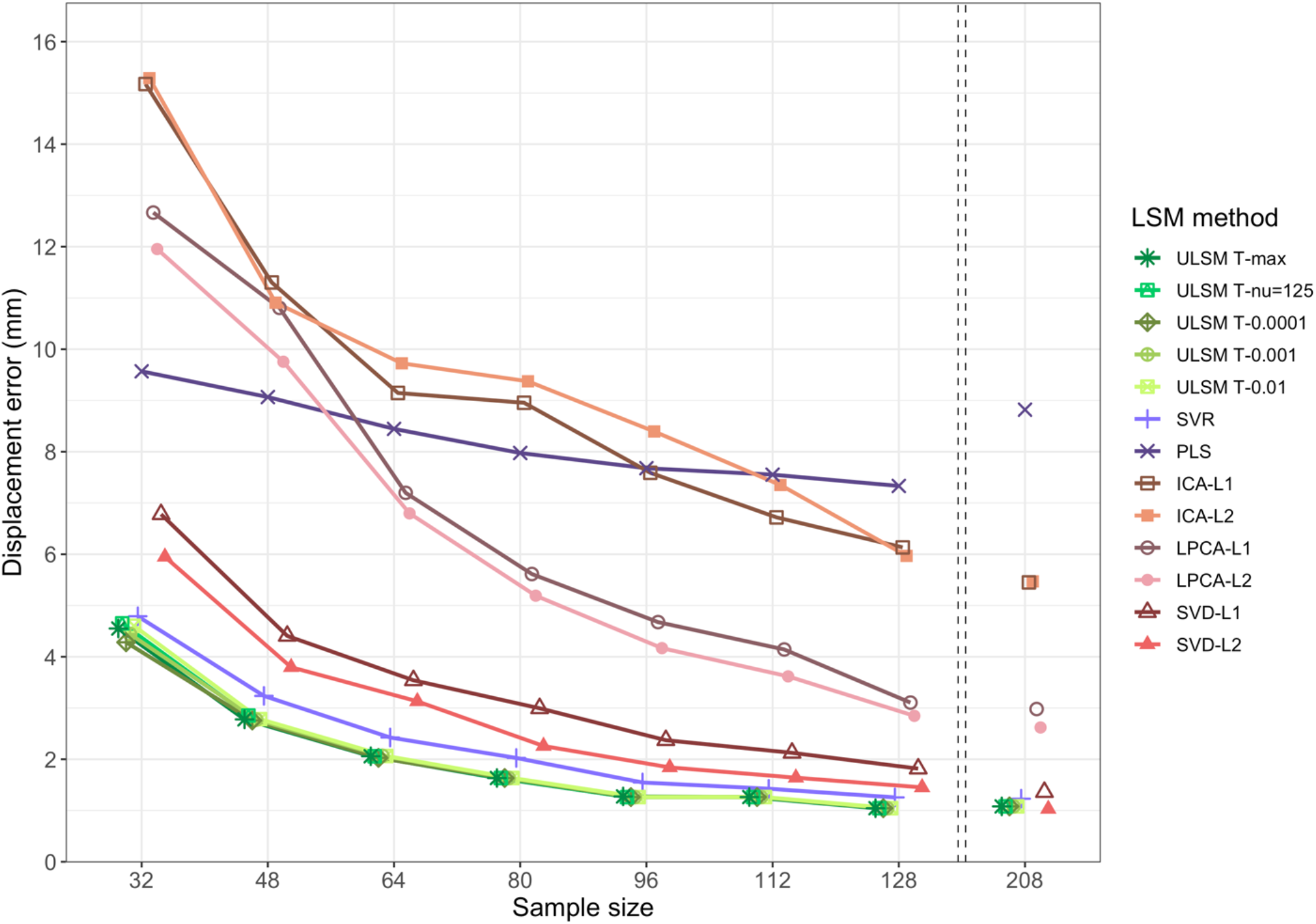
Displacement (in mm) of LSM output map position for single target simulations across different LSM methods at different sample sizes calculated as the average distance between maximum statistic location (Max_LSM_) and nearest location on the target parcel to the LSM output map (Closest_target_). The left side of this figure focuses on small sample sizes as most representative of typical LSM studies, since minimal improvements in accuracy are observed for samples larger than 128.

**Figure 7.**
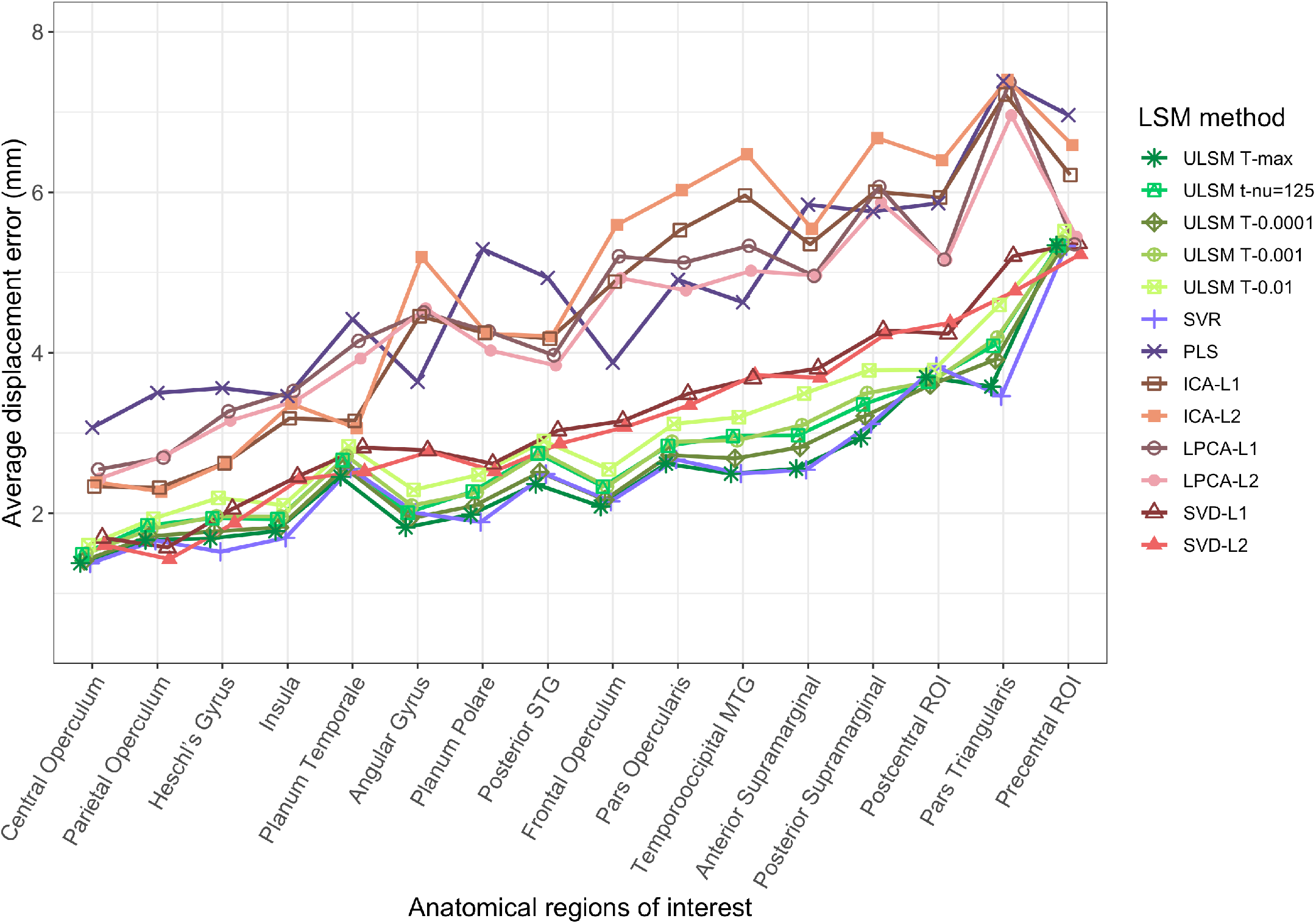
Average displacement error (in mm) of LSM output map position for single target simulations across different anatomical parcels and LSM methods.

**Figure 8.**
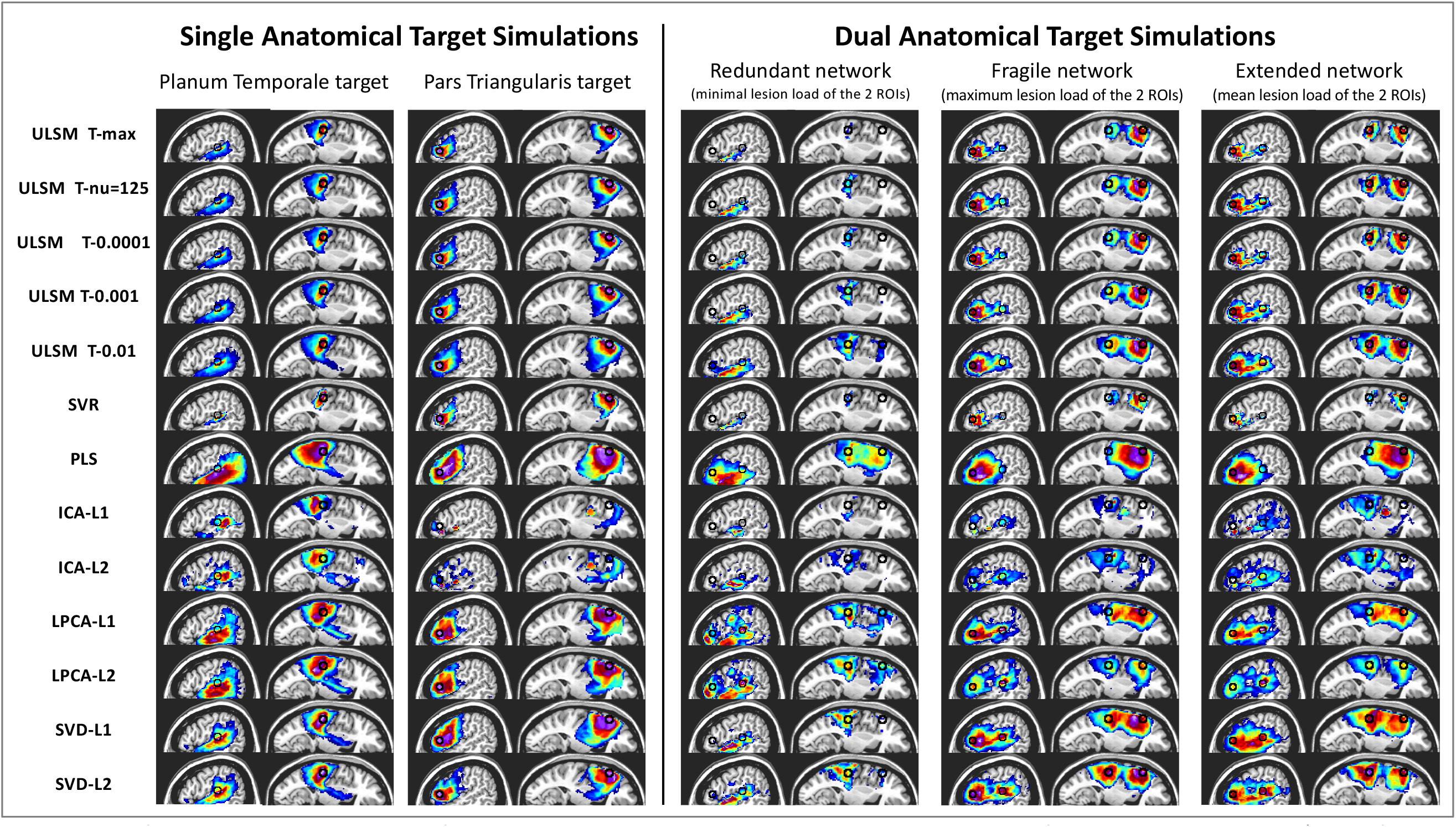
Left side: LSM output maps for single target simulations across all LSM methods for Planum Temporale (one of the most accurately detected regions) and Pars Triangularis (one of the least accurately detected regions) for a sample size of 64 (typical for single target simulations), medium behavioral noise level with lesion mask smoothing (4 mm). Right side: LSM output maps for dual-target simulations for these two regions for the three network types for a sample size of 112, medium behavioral noise level, and lesion mask smoothing (4 mm). On all the LSM maps the location of the target is denoted by a black circle placed at the center of mass of the corresponding anatomical parcel(s) generating the synthetic score. The circle is used for visualization purposes only.

As expected, the average spatial displacement error varied significantly depending on which metric was used for determining LSM output map location (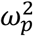 = .62; COM_LSM,_ wCOM_LSM,_ Max_LSM_) or target location (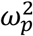 = .82; COM_target_, Closest_target_). Use of Max_LSM_ as the measure of LSM output map and Closest_target_ as the measure of target parcel location led to the highest accuracy estimates. Further, different LSM methods demonstrated varying levels of accuracy, depending on the chosen measure of LSM output map location (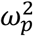 = .56) and target location (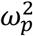= .36). In general, MLSM methods showed greater disparity between COM_LSM_ and wCOM_LSM_ measures, while ULSM methods demonstrated a greater difference between wCOM_LSM_ and Max_LSM_ measures, with the COM_LSM_ and wCOM_LSM_ measures yielding more similar findings. More dense LSM solutions showed greater disparity between COM_target_ and Closest_target_ measures. Additionally, measures of LSM output map location strongly interacted with Behavioral Noise Level (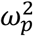 = .81), Sample Size (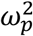 = .81), and Mask Smoothing (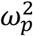 = .54). With higher behavioral noise levels, mean centroid location measurements (both COM_LSM_ and wCOM_LSM_) improved, while accuracy measured via Max_LSM_ decreased. wCOM_LSM_ was the most stable across all noise levels. With larger sample size, spatial accuracy improved, particularly, as indexed by the Max_LSM_ measure. Lesion mask smoothing was most beneficial for the Max_LSM_ measure. Tables with the mean simulation values for these analyses can be found in Appendix A (Tables A1-A3).

An additional post-hoc analysis was run with smaller-sized anatomical targets (n=30) and the highest behavioral noise level (0.71 SD), with and without lesion size as a covariate. Using lesion size as a covariate resulted in higher spatial accuracy of LSM output maps across all sample sizes and all LSM methods (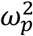 = .87, mean displacement of 8.6 mm vs. 12.1 mm without it). The effect of covarying for lesion size varied as factor of LSM Method (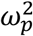 = .43) with MLSM methods showing more improvement in spatial accuracy (3 – 8 mm) relative to ULSM methods (2 – 3 mm) with the lesion size covariate (see Appendix A, Table A4).

#### 3.1.3 Spatial accuracy with overlap-based metrics

Spatial accuracy as measured by dice coefficient values varied significantly across LSM Methods (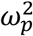 = .93). According to this measure ULSM (T-max and T-nu=125) and SVR had the highest spatial accuracy. However, the mean dice index values were very low across all LSM methods (ranging from .17 to .02; see Table 1), rendering the dice coefficient relatively uninformative as a metric for evaluating overlap between LSM output map and the target anatomical parcel. Behavioral Noise Level (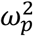 = .86), Sample Size (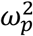 = .79), and Mask Smoothing (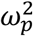 = .41) all affected the spatial accuracy as measured by dice coefficients. Generally, dice coefficients were larger (more accurate) with increased behavioral noise levels and no mask smoothing but decreased with larger sample sizes as the significant clusters become larger and extended beyond the target parcel. First order interactions between LSM Method x Sample Size (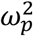 = .49) and LSM Method x Behavioral Noise Level (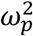 = .46) were significant. Variations in the dice coefficient as a function of sample size and behavioral noise level across different LSM methods are presented in Figure 9. As can be seen in this figure, ULSM methods and SVR showed steeper decreases in dice coefficient values as sample size increased.

**Figure 9.**
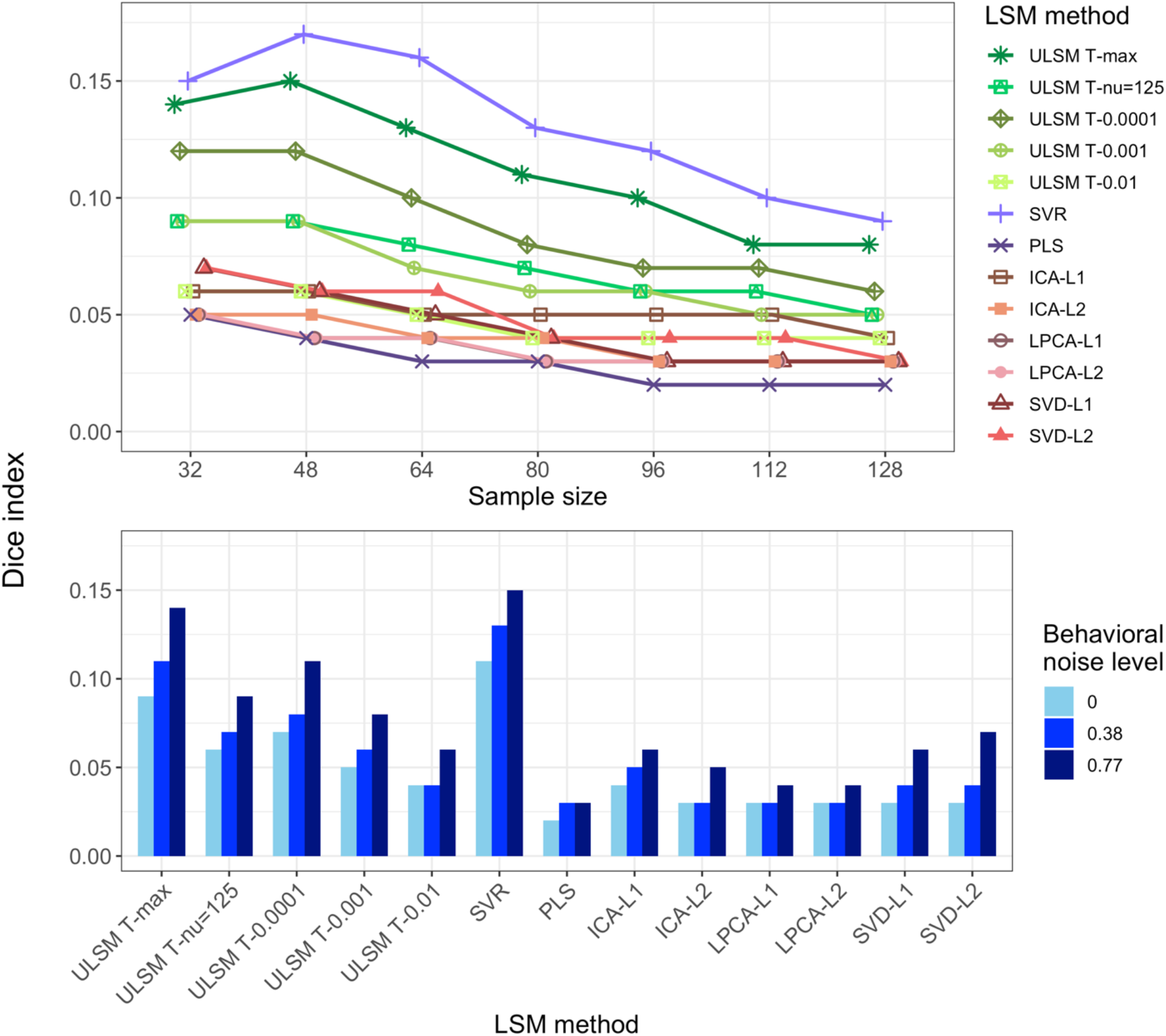
Dice index as a function of LSM method and sample size (top panel) or behavioral noise level (bottom panel) for single target simulations.

In contrast to the dice coefficient values, OSK distribution values were substantially higher and showed higher variability across factor levels, rendering it more useful than the dice coefficient for evaluation of spatial accuracy (see Table 1). The OSK distribution statistic greatly varied across different LSM methods (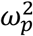 = .85). Liberal ULSM methods (ULSM T-0.01 and T-0.001) and SVD showed the highest OSK values, with the advantage being particularly evident with smaller sample sizes. Behavioral Noise Level (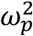 = .96), Sample Size (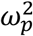 = .95), and Mask Smoothing (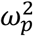 = .80) all had a strong effect on the OSK statistic. Larger sample sizes, lower noise levels and mask smoothing resulted in higher OSK distribution statistic values. First order interactions between these factors also demonstrated moderate effect sizes (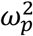 ranged from .36 to .50). LSM methods showed varying degrees of improvement as sample size increased (Figure 10, top panel). ICA-L1, ICA-L2, SVR, and ULSM T-max performed the worst at small sample sizes but showed the most improvement with increased sample sizes, although still performing below the remaining LSM methods. Higher behavioral noise levels required larger sample sizes to achieve similar levels of accuracy (Figure 10, middle panel). Further, different LSM methods demonstrated variable susceptibility to behavioral noise, with sparser solutions showing higher susceptibility to behavioral noise (more pronounced decrease in OSK values) (Figure 10, bottom panel).

**Figure 10.**
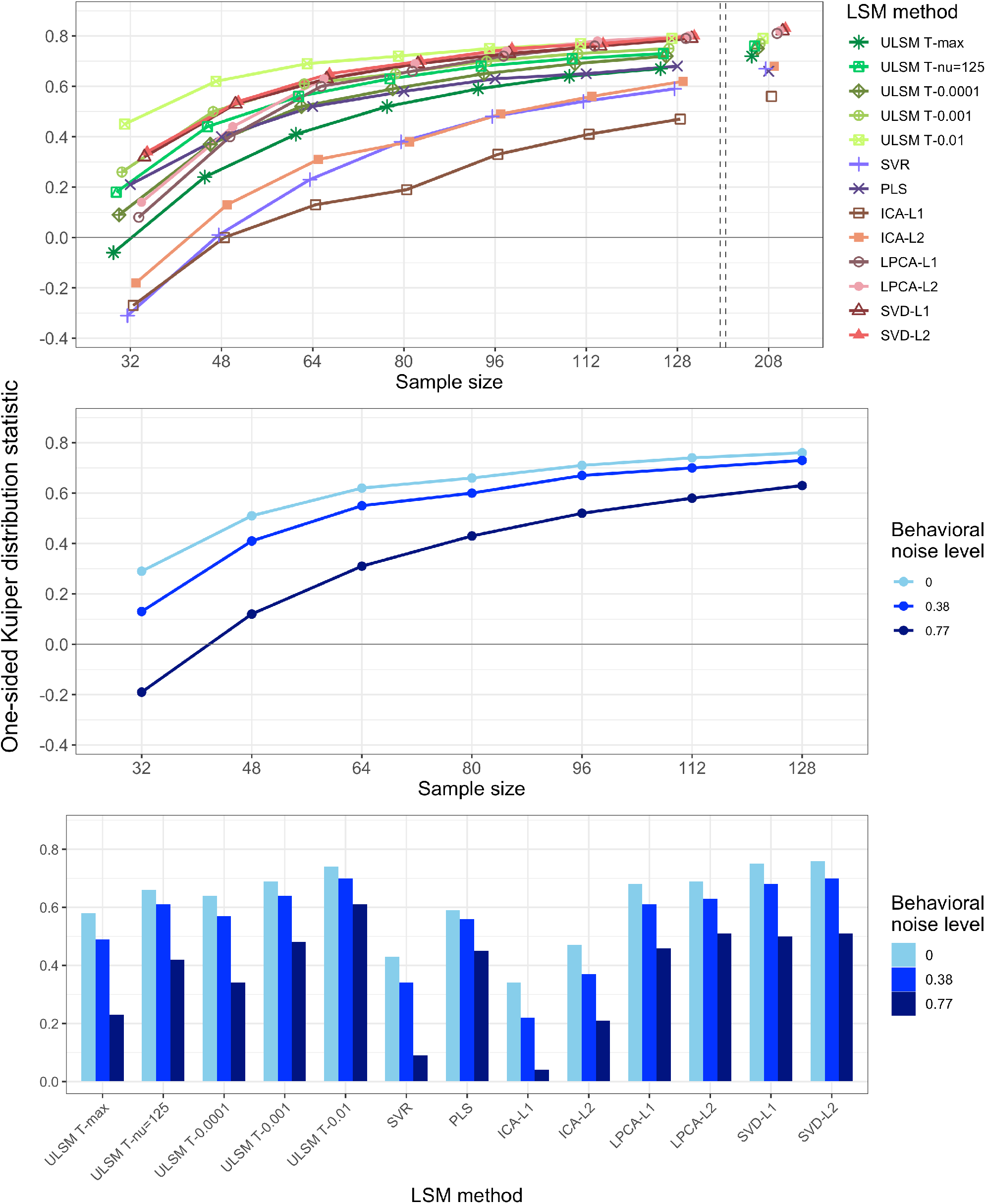
One-sided Kuiper (OSK) distribution statistic as a function of the LSM method and sample size (top panel), behavioral noise level and sample size (middle panel) or LSM method (bottom panel) for single target simulations.

### 3.2 Dual anatomical target simulations

Dual-target simulations were run on three network types: Redundant, Fragile, and Extended. LSM Method, Sample Size and Type of Network all had a strong effect on power (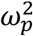 ranged from .80 to .86; see Table 2 for mean simulation values). A sample size of 64 or larger was required to achieve 80% power for all three network types. Two-way interactions between these factors demonstrated medium to large effect size (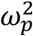 ranged from .36 to .50) (see Appendix A, Tables A5-A6 for relevant data). The Redundant network was the hardest to detect for all LSM methods, requiring a much larger sample size to obtain a level of power similar to that of the Fragile and Extended networks.

**Table 2.** Mean simulation values for all manipulated factors for dual anatomical target

| LSM Method | ULSM<br>T-max | ULSM<br>T-nu=125 | ULSM<br>T-0.0001 | ULSM<br>T-0.001 | ULSM<br>T-0.01 | SVR | PLS | ICA-L1 | ICA-L2 | LPCA-L1 | LPCA-L2 | SVD-L1 | SVD-L2 |
| --- | --- | --- | --- | --- | --- | --- | --- | --- | --- | --- | --- | --- | --- |
| Power (%) | 98.6 | -* | 98.7 | 99 | 99.1 | 98.5 | 96.7 | 97.4 | 98.4 | 98.2 | 98.6 | 95.9 | 95.7 |
| Dice index | 0.13 | 0.11 | 0.12 | 0.11 | 0.09 | 0.14 | 0.05 | 0.07 | 0.06 | 0.06 | 0.07 | 0.07 | 0.07 |
| OSK statistic | -0.21 | -0.05 | -0.12 | 0 | 0.13 | -0.34 | 0.02 | -0.2 | 0 | 0.2 | 0.25 | 0.26 | 0.25 |

| Sample size | 64 | 80 | 96 | 112 | 128 | 144 | 160 | 176 | 192 | 208 |
| --- | --- | --- | --- | --- | --- | --- | --- | --- | --- | --- |
| Power (%) | 92.8 | 95.8 | 97.6 | 98.2 | 98.8 | 99.1 | 99.3 | 99.7 | 99.7 | 99.8 |
| Dice index | 0.1 | 0.1 | 0.09 | 0.09 | 0.09 | 0.09 | 0.08 | 0.08 | 0.08 | 0.08 |
| OSK statistic | -0.27 | -0.17 | -0.09 | -0.02 | 0.03 | 0.08 | 0.11 | 0.13 | 0.16 | 0.18 |

| Type of Network | Redun-<br>dant | Fragile | Exten-<br>ded |
| --- | --- | --- | --- |
| Power (%) | 94.7 | 99.7 | 99.8 |
| Dice index | 0.09 | 0.09 | 0.09 |
| OSK statistic | -0.11 | 0.04 | 0.12 |

| Anatomical Parcels | Mean | SD | Max | Min | Quart<br>1 | Quart<br>3 |
| --- | --- | --- | --- | --- | --- | --- |
| Power (%) | 98.1 | 0.1 | 99.5 | 96.4 | 97.4 | 98.8 |
| Dice index | 0.09 | 0.02 | 0.12 | 0.06 | 0.07 | 0.11 |
| OSK statistic | 0.01 | 0.2 | 0.22 | -0.46 | -0.01 | 0.11 |

Note. \* - Power not calculated due to a technical error.

Spatial accuracy of the detected networks (as determined by the OSK overlap measure) greatly varied based on the LSM Method (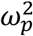 = .95). SVD and LPCA (both L1 & L2), as well as ULSM T-0.01 showed consistently positive values (implying that values inside the target parcels were higher than outside) for sample size greater than 80 (see Figure 11, top panel). Overall, OSK values were much lower for the dual-target simulations. All LSM methods were numerically less accurate in detecting anatomical networks compared to the single target situation (average OSK across all LSM methods of 0.01 for networks vs. 0.5 for single targets; for best performing MLSM method OSK of 0.25 vs 0.66 respectively). Effect size for Sample Size (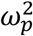 = .98), Type of Network (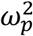 = .84), and first order interactions of LSM Method x Type of Network (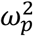 = .54) and LSM Method x Sample Size (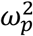 = .61) were also large. The Extended network resulted in the highest spatial accuracy, and the Redundant network resulted in the lowest spatial accuracy across all LSM methods (se Figure 11, bottom panel). A similar pattern was observed for the dice coefficient values, although with significantly smaller values and smaller variation between LSM methods (see Table A7 in the Appendix). See Figure 8 for LSM output maps for the three network types for planum temporale and pars triangularis regions across different LSM methods.

**Figure 11.**
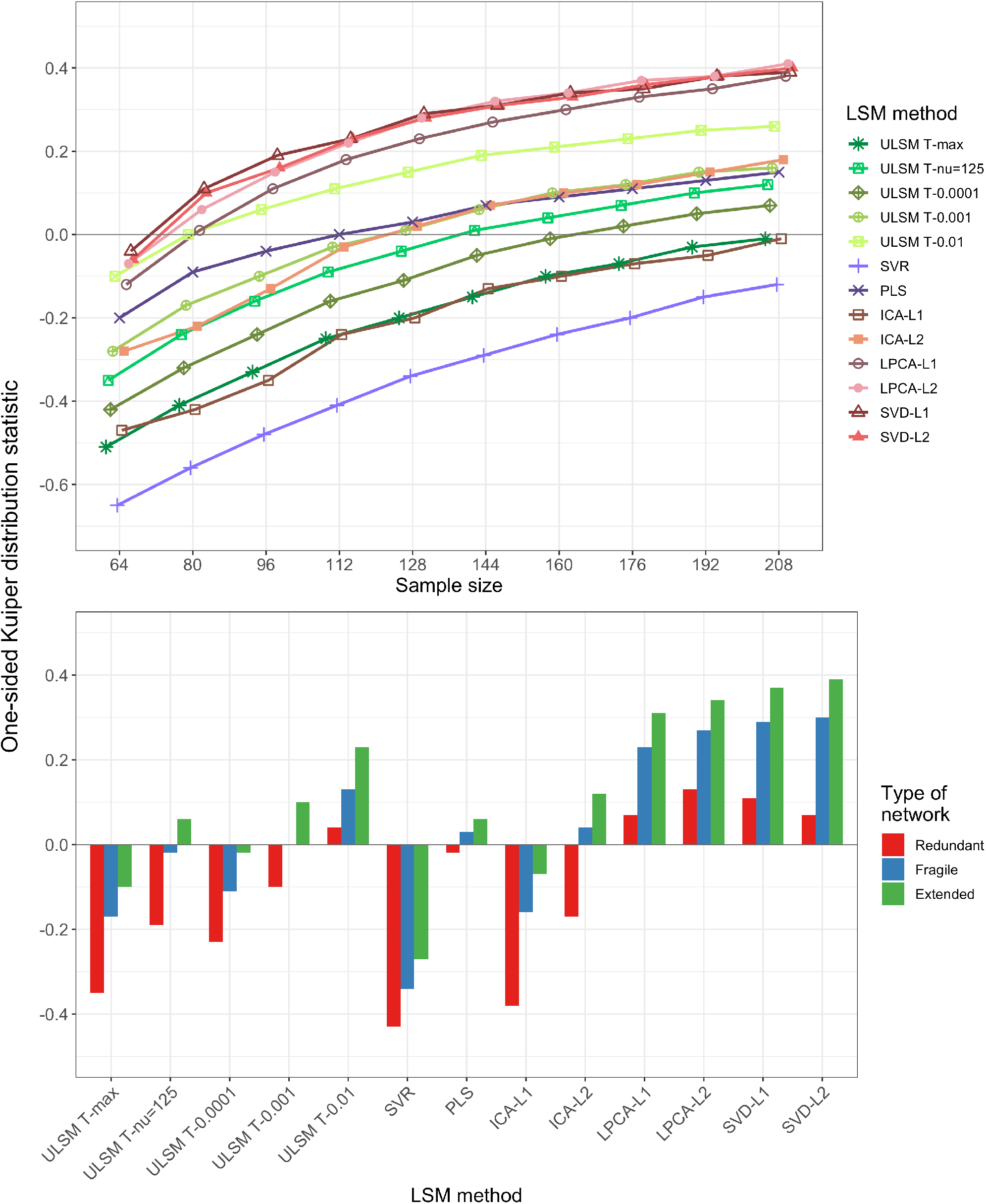
One-sided Kuiper (OSK) weighted-overlap statistic as a function of the LSM method and sample size (top panel) or type of network (bottom panel) for dual-target simulations.

### 3.3 False positive simulations

For the false positive simulations, SVR and ULSM T-max performed best, yielding the smallest number of clusters with the smallest number of voxels (see Table 3). Sample Size and Mask Smoothing did not have a significant impact on the rate of false positives. A Pearson correlation analysis showed that false positive rates were not consistent across different LSM methods. The correlation of false positives was lowest between ULSM and MLSM methods (see Figure 12). In particular, PLS and SVD had low correlations with the other methods.

**Figure 12.**
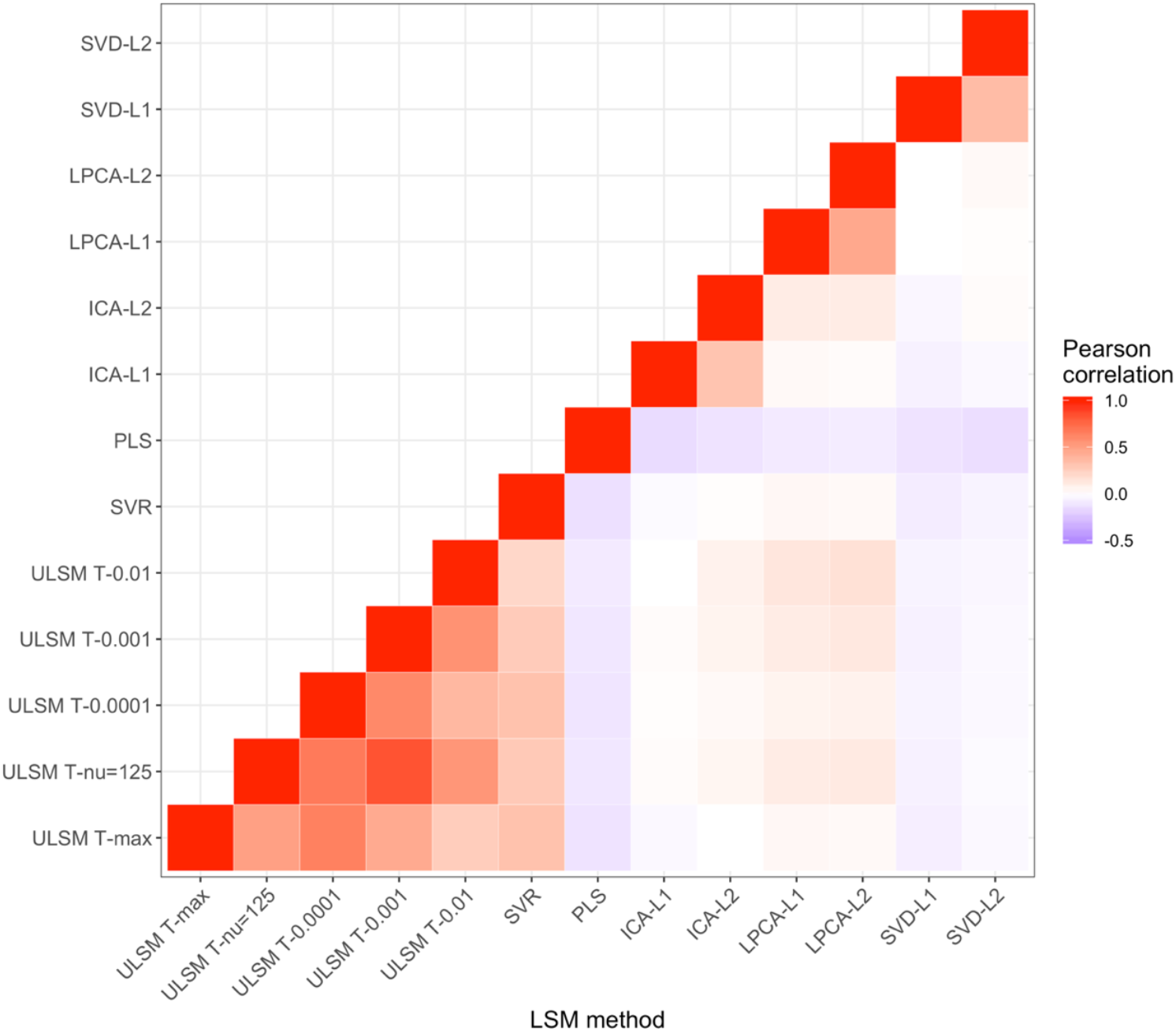
Pearson correlations between false positive rates across different LSM methods.

**Table 3.** Mean values of above-threshold cluster characteristics.

|  | Number<br>of clusters | Total number<br>of voxels | Maximum<br>statistic |
| --- | --- | --- | --- |
| **T-max** | 1.5 | 17 | 4.8 |
| **T-nu=125** | 4.6 | 312 | 4.6 |
| **T-0.0001** | 1.2 | 79 | 4.7 |
| **T-0.001** | 1.0 | 473 | 4.5 |
| **T-0.01** | 1.0 | 2429 | 4.4 |
| **SVR** | 1.5 | 17 | - |
| **PLS** | 5.5 | 2706 | - |
| **ICA-L1** | 4.7 | 644 | - |
| **ICA-L2** | 5.3 | 712 | - |
| **LPCA-L1** | 5.4 | 1603 | - |
| **LPCA-L2** | 5.2 | 1957 | - |
| **SVD-L1** | 7.1 | 903 | - |
| **SVD-L2** | 7.5 | 982 | - |

*Note.* Maximum statistic is only interpretable for ULSM methods, since it is scaled for MLSM methods (always between 0 and 1).

### 3.4 LSM analysis of real behavioral data

We next compared results across the 13 different LSM methods, using real behavioral data collected from 168 patients from the NorCal dataset. The behavioral data consisted of WAB language subscores for speech fluency, single-word auditory comprehension, and verbal repetition (Kertesz,1982, 2007).

Results showed distinct areas identified for different behavioral scores, such as frontal regions for speech fluency and posterior temporal regions for single word comprehension (see Figure 13 for LSM maps across methods). However, as can be seen in Figure 13, substantial variability in the identified regions was observed between different LSM methods. MLSM methods tended to identify more regions, including regions not typically associated with the behavior (e.g., frontal regions for single-word comprehension). In contrast, ULSM maps typically clustered around the maximum statistic location more traditionally associated with the behavior (e.g., posterior temporal cortex for single-word comprehension). The methods that provided more sparse solutions (smaller LSM clusters) showed better differentiation between different behaviors under examination (e.g., conservative ULSM methods and ICA).

**Figure 13.**
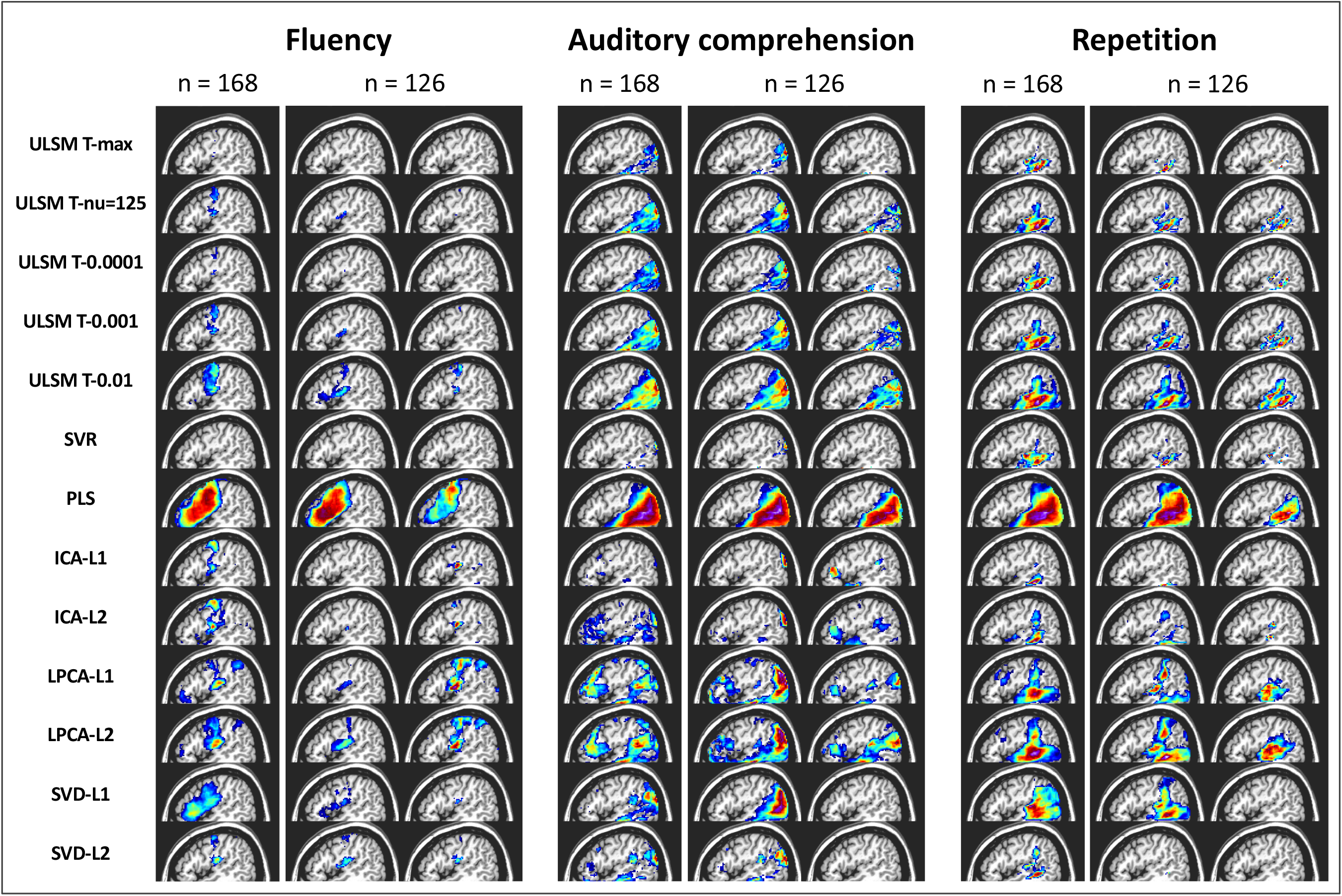
LSM maps by method for three language variables (speech fluency, single-word auditory comprehension, verbal repetition): for the full sample (n = 168) and two random subsamples (n = 126).

Since there is no ground truth in the case of real behavioral data, we evaluated accuracy in terms of the stability of the obtained solutions using a random subsampling approach for n = 126 (using the full sample as the target map). The LSM maps produced in the subsample analysis varied between themselves and often clearly differed from the full sample map (see again Figure 13). Stability varied substantially across LSM techniques, as shown by spatial metrics averaged across the 100 subsample analyses and by the average correlation between the full LSM output and each of the subsample analyses (see Table 4). The methods that provided more dense solutions (larger LSM clusters) tended to show higher stability of results (e.g., liberal ULSM methods and PLS). Moderately conservative ULSM methods showed higher stability of results compared to MLSM as indicated by lower average displacement and higher OSK statistic, as well as higher correlations.

**Table 4.**
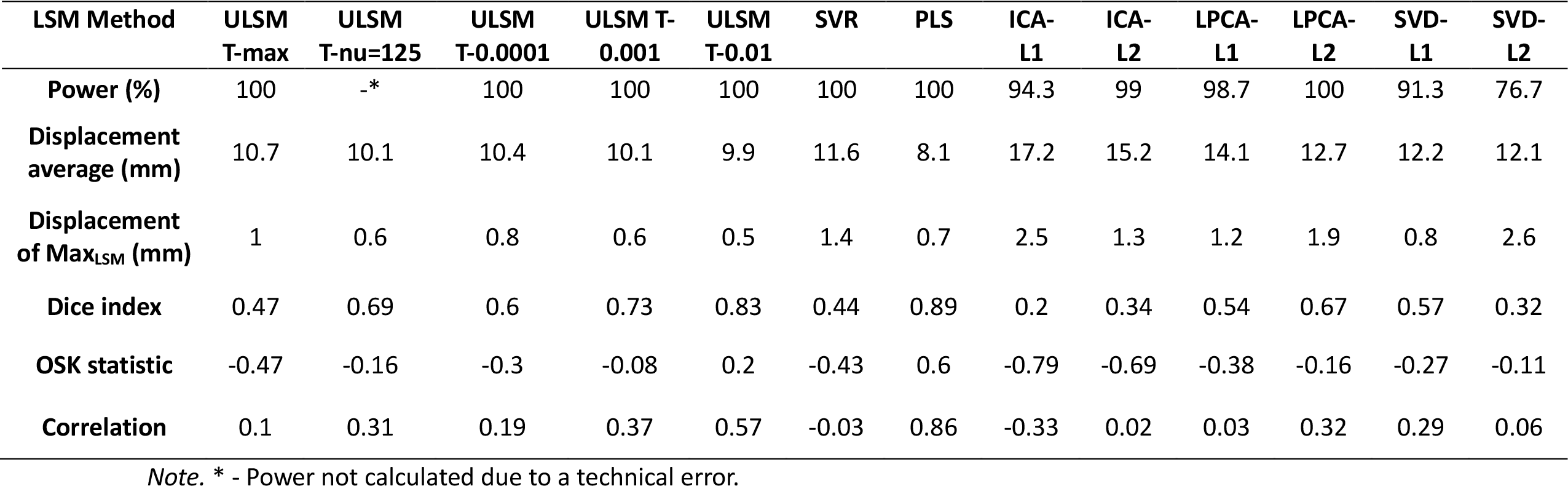
Mean subsampling analyses results: percentage of trials where significant voxels were detected (Power (%)), average displacement across all distance metrics (Displacement average(mm)), average distance between Maximum statistic and any voxel within the target ROI (Displacement of Max_LSM_ (mm)), Dice index, One-sided Kuiper distribution statistic (OSK statistic: −1 worst to +1 best), and correlations between the full sample and the subsampling LSM maps (Correlation).

Finally, when evaluating the stability of the subsampling analyses substantial variability across different distance-based metrics was observed (see Table 5). The maximum LSM statistic of each subset LSM map usually landed on or very near the full target LSM map (0.3 mm), but often not very close to the full map’s maximum value (20.6 mm). Since the location of the maximum statistic is most commonly reported in LSM papers, we also compared the stability of the 13 different LSM methods using this metric (based on mean Max_LSM_ to Closest_target_; see Displacement of Max_LSM_ in Table 4). Again, moderately conservative ULSM methods showed higher stability (range: 0.6 – 0.8 mm) relative to MLSM methods (range: 0.8 – 2.6 mm) with the exception of PLS.

**Table 5.** Mean accuracy of LSM subsampling analyses.

|  |  | Target parcel map position |  |  |  |
| --- | --- | --- | --- | --- | --- |
|  |  | COM | wCOM | Max | AnyHit |
| LSM<br>output<br>map<br>position | COM | 7.9 | 7.7 | 20.6 | 1.7 |
|  | wCOM | 8.2 | 7.8 | 20.2 | 1.6 |
|  | Max | 21.2 | 20.1 | 20.6 | 0.3 |

## 5 DISCUSSION

In the current study, we conducted the first comprehensive, empirical comparison of several univariate and multivariate LSM methods. Statistical power, spatial accuracy and robustness of each method was evaluated using both synthetic and real datasets across a range of relevant parameters, including sample size, lesion smoothing, noise level, and type of network. A number of different metrics were used to evaluate spatial accuracy. Cumulatively, our analyses indicated that both ULSM and MLSM methods can be equally robust in locating brain behavior relationships, depending on the design of the study, the research question being asked, and under a reasonable interpretation. The results provide crucial insights into the accuracy of different LSM methods and their susceptibility to artifact, providing a first of its kind data-driven navigational guide for users of LSM analyses.

### 5.1 Comparison of ULSM and MLSM methods

For single anatomical target simulations with synthetic behavioral data and real lesion data, ULSM methods with conservative FWER thresholds (e.g., T-max and T-nu=125) and some of the simpler data reduction (e.g., SVD-based) and regression-based (particularly SVR) MLSM methods showed comparably good spatial accuracy. The majority of MLSM methods required on average 10-20 more participants than ULSM methods, in order to achieve a similar level of statistical power and spatial accuracy.

With respect to dual-target simulations (i.e., identifying a network), ULSM methods with more liberal cluster-based FWER thresholds (e.g., T-0.01) and some of the MLSM data reduction methods (e.g., LPCA) demonstrated the highest spatial accuracy. The dice coefficient values for dual-target simulations were low across all LSM methods and network locations. Thus, if a dice coefficient is used as a measure of accuracy (as was done previously in Pustina et al., 2018), there is no practical difference between ULSM and MLSM methods in their ability to detect and localize networks (cf. the dice values in Pustina et al.). However, if LSM statistical values are taken into account when interpreting localization of results (e.g., the OSK distribution of inside target vs. outside target statistics), there are advantages to some of the MLSM methods (LPCA, SVD) in detecting dual-target networks. Still, the accuracy of localization of dual-target networks was substantially lower overall than localization of a single target across all methods. Thus, all LSM methods, both ULSM and MLSM show a limited ability to accurately identify dual-target networks. Further, our results unequivocally highlight the importance of having a sample with ≥ 100 participants in order to have sufficient power and accuracy to detect dual-target networks, in particular, redundant networks, which were more difficult to accurately identify across methods.

Additionally, our results clearly demonstrate that not all MLSM methods are equally good at detecting networks. For instance, SVR was comparable to the more conservative ULSM methods in the current study (see Sperber et al., 2019 for more on this). In general, SVR performed similarly to ULSM methods across all simulations, highlighting the fact that the version of SVR we used with default settings (DeMarco & Turkeltaub, 2018) was tuned to work more like a ULSM method.

With respect to false positives, ULSM methods with conservative FWER thresholds (T-max and T-nu=125) and regression-based MLSM methods performed best. Specifically, ULSM T-max and MLSM SVR had the lowest false positive rates. Most of the MLSM methods generally separated the false positive solutions into a large number of spatially disconnected clusters (i.e., false positive networks). Thus, these false positives parallel the kinds of solutions that each method is optimized to detect when there is a valid brain-behavior relationship. Not surprisingly, there were moderate correlations in false positive clusters between the 5 ULSM variants and SVR and also between similar data reduction techniques (primarily L1 and L2 solution types). In contrast, there were low correlations between false positive clusters in ULSM versus MLSM methods. Based on these findings, we suggest that both ULSM and MLSM methods should be used in tandem, given an adequate sample size for each method, to increase confidence in the results of an LSM analysis. Real findings will be consistent across different classes of LSM methods, but spurious findings will be inconsistent.

With respect to LSM performance on the real behavioral data, moderately conservative ULSM (T-nu=125 and T-0.001) and specific MLSM methods (ICA, SVD) identified largely similar regions. This is again in line with previous empirical studies that employed both ULSM and MLSM methods to investigate brain-behavior relationships and yielded highly coherent results between the two approaches (e.g., Fridriksson et al., 2018; Thye & Mirman, 2018). However, some MLSM methods also identified brain regions that are not typically associated with the behavior under examination. Also, stability of the obtained solutions was generally higher for ULSM methods. Specifically, moderately conservative ULSM methods (T-nu=125 and T-0.001) provided a good balance between a sparse, spatially differentiated solution and robustness (stability) of obtained results. Methods that provided more dense solutions tended to show higher stability of results (e.g., PLS, ULSM T-0.01). However, these methods showed less spatial discrimination with results obtained for different language functions, yielding more similar LSM output maps relative to the sparse solutions. The opposite pattern was observed for methods providing sparse solutions (e.g., ULSM T-max, SVR, SVD-L2, ICA-L1): While they provided clearly distinguishable solutions for the various language functions, these solutions were not as robust. Interestingly, although in general SVR performed similarly to ULSM methods, the stability of LSM maps obtained with SVR was lower and more akin to MLSM methods.

Taken together, these results provide crucial insights into application of different LSM methods. While no single method produced a thresholded LSM map that even approached an exclusive delineation of brain regions associated with the target behavior, under specific conditions certain methods performed better than others. ULSM methods demonstrated particularly good performance in terms of accuracy and robustness for identifying single targets. Here MLSM methods clearly lagged in power, accuracy, susceptibility to behavioral noise, and stability. Although, not all MLSM methods were equally good at identifying networks, in general, they performed better than ULSM. This underscores the notion that no single LSM method can provide an ultimate solution for establishing brain correlates of cognitive functioning.

### 5.2 Impact of other parameters on LSM accuracy

In addition to the comparison of different LSM methods, we also investigated the impact of a number of other parameters on mapping accuracy across methods, including anatomical target location and size, mask smoothing, behavioral noise level, sample size and lesion volume. With respect to target location, we found variable accuracy across spatial locations, with especially poor performance in cortical locations on the edge of the lesion masks (areas of lower power). The size of target parcels (small or large) did not have a substantial impact on the LSM results, with accuracy being only slightly higher for the smaller parcels in the single anatomical target simulations. This implies that the current findings are applicable to LSM analyses irrespective of the size of the hypothetical target parcel(s).

We also compared the effects of lesion mask smoothing at 0 versus 4mm Gaussian FWHM. The 4mm smoothing led to improved accuracy of localization across all LSM methods. We suspect that with real data, smoothing should be even more beneficial given that in simulations with synthetic behavioral scores, there was effectively no lesion delineation error, anatomical normalization error, or anatomical size/shape mismatch across patients as there ordinarily is (typically ∼5mm in extent) in stroke patients (Crinion et al., 2007; Rorden et al., 2012).

Behavioral noise level was also manipulated in the current study, in order to investigate its impact on accuracy of localization. More behavioral noise in small sample sizes (n < 100), led to a reduction in statistical power, particularly for MLSM methods. At larger sample sizes (n > 100), behavioral noise did not considerably diminish power. Behavioral noise also negatively impacted the accuracy of mapping, as evidenced by reduced OSK values, but had no noticeable impact on distance-based measures of spatial accuracy.

With respect to sample size, the current findings showed improved power and accuracy with increasing numbers of patients, not surprisingly. It is important to note, however, that the benefit of increased sample sizes on spatial accuracy plateaued at about 130 participants for all LSM methods in single target simulations. LSM methods do not appear to be spatially unbiased statistical methods, and stroke lesions are not entirely random (Sperber & Karnath, 2016), so beyond a certain sample size the increase in coverage and in variability of lesion patterns is minimal leading to muted effects on accuracy.

Finally, including lesion volume as a covariate significantly improved spatial accuracy across all LSM methods. The importance of including lesion volume as a covariate has been shown in previous studies for ULSM (Sperber & Karnath, 2017) and some MLSM methods (DeMarco & Turkeltaub, 2018). Our current findings further reinforce the stipulation that a lesion volume covariate should be included in all types of LSM analyses.

### 5.3 Different metrics of LSM accuracy

Overall, our simulated results show that the Max_LSM_ measure (location of the maximum statistic of the LSM output map) is the most accurate metric for identifying the location of the LSM cluster. Similarly, Pustina et al. (2018) also demonstrated that peak voxel displacement measure provided the most accurate mapping for both ULSM and MLSM methods. The utility of using Max_LSM_ is reinforced by our results with real behavioral data, as this metric demonstrated the smallest displacement error relative to other measures in the subsampling analysis. At the same time, the wCOM_LSM_ value (mean centroid location weighted by statistical values of the LSM output map) was the most robust metric across varying behavioral noise levels.

In our study, the dice index metric was very low across all LSM methods and across all spatial locations for both single and dual-target simulations (similar to Pustina et al., 2018). In contrast, the OSK distribution comparison statistic takes into account both statistical values and their localization, and thus can be considered a type of weighted dice index. Accordingly, when the statistical values were taken into account both inside and outside the target location, a more nuanced picture emerges, highlighting advantages for certain methods (e.g., ULSM methods with liberal cluster-based FWER thresholds, SVD, LPCA) optimized for more dense solutions. Good OSK performance implies that higher statistical values end up inside the target ROI. This further reinforces the idea that statistical values should be taken into account when mapping LSM cluster location, rather than treating all above threshold locations equally and uniformly localizing the whole cluster.

### 5.4 Limitations and directions for future work

Similar to previous simulation studies, we modeled a linear relationship between impairment and lesion location. In reality, brain-behavior relationships probably follow a more complex pattern and the presented simulations might not be fully representative of real behavioral data in the chronic stages of stroke. Furthermore, there were no location-independent lesion size behavioral effects included in the simulations with synthetic behavioral data. However, there is no current consensus about what these more complex patterns would be and, thus, it is unclear how to model them. Additionally, instead of linear regression, other statistics could be employed in the ULSM analyses. In the current study we used linear regression rather than logistic regression (appropriate when the dependent variable is bounded) because linear regression is much faster to compute, is more familiar to users, and is standard in previous studies (Akinina et al., 2019; Baldo et al., 2013, 2018; Ivanova et al., 2018). While linear regression is sufficient to model the perfectly linear relationships in synthetic data, real behavioral data might indicate a different approach such as the use of nonparametric measures (Rorden et al., 2007).

Further, efficiency of control factors beyond lesion volume was not investigated. Future work should explore the impact of additional control factors, such as the “anatomical bias” vectors (Sperber & Karnath, 2017), on accuracy of mapping of different LSM methods. Their implementation can potentially further improve spatial accuracy of (specific) LSM methods.

Another limitation of the current study is the lack of a sparse version of PLS or similar methods (like SCCAN, Pustina et al., 2018). We did not implement SCCAN in the current study because it is highly computationally intensive given that one needs to determine several hyperparameters in order to optimize performance for a given hypothesis/target configuration. Potentially, it might be superior to the MLSM methods evaluated in the current study. However, the results compiled in Pustina et al. (2018) – low dice scores for outputs (particularly for dual-target simulations) but good performance for maximum distance measures (similar to ULSM methods) – is consistent with our basic findings here. It is likely that the decision by SCCAN creators to allow tuning of hyperparameters will give it an advantage over the current implementation of SVR, where the hyperparameters were fixed to reproduce ULSM output. In fact, we predict that if SVR were implemented with the same cross validation hyperparameter tuning that SCCAN uses, it might match or exceed the performance of MLSM methods tested above in dual-target simulations (at the cost of extra computational time and perhaps an increase in sample size). Alternatively, it would be important to investigate if SCCAN would perform as well with larger target ROIs, since originally it has been tuned to detect sparse solutions, and in Pustina et al., 2018 very small parcels were used as targets. Also, the robustness of SCCAN’s solutions needs to be determined. All these outstanding issues require further investigation along with a more in-depth and systematic exploration of how tuning of features ((hyper)parameters; e.g., cost, gamma, sparseness) might affect the results different MLSM methods produce.

### 5.5 Recommendations for LSM analysis

In summary, our simulations show that both ULSM and MLSM methods are similarly effective with respect to the majority of brain mapping analyses. The choice of a particular LSM method is largely dependent on the goals of a specific study. When there is a hypothesis specific to a single anatomical target, ULSM methods with appropriate corrections (e.g., lesion volume and permutation-based correction for multiple comparisons) are most appropriate, as ULSM methods are generally more accurate and robust at detecting and localizing a single target with a low false positive rate. When a study is predicting a dual (or multi) target network, certain MLSM methods (LPCA-L2 and SVD-L2) and ULSM methods with liberal cluster-based thresholds (ULSM T-0.01) are superior for accurately identifying multiple anatomical targets in a network. For studies with small sample sizes (n = 50-80), ULSM methods are preferred, as a majority of MLSM methods (except SVR) require a larger number of participants to achieve a comparable level of accuracy.

There are also a number of recommendations that extend from the current findings which apply to all LSM techniques. First, it is imperative to have sufficient statistical power, not just for the usual reason of minimizing false negatives, but because sufficient power in LSM analyses decreases the spatial error in localization. It is recommended that power analyses be performed for each analysis and that studies include a minimum of 50 participants with moderate noise level in the data (as can be typically expected with behavioral data) to obtain reliable results. Also, if one is hypothesizing a network of regions, we recommend a sample size of *n* ≥ 100. Studies with small sample sizes (< 50) should utilize alternate types of lesion-based analyses (e.g., lesion overlays), as no LSM method is reliable in such instances.

Our results also show that lesion mask smoothing at 4mm improves accuracy of localization across all metrics, even in situations with no misalignment or lesion mask drawing errors (i.e., synthetic data). Thus, we recommend mask smoothing to diffuse possible bias in lesion segmentation and normalization. Lesion volume should always be included as a covariate for all LSM methods, as it mitigates some of the spatial autocorrelation effects and results in significant improvement in accuracy of localization across all sample sizes and LSM methods.

The final recommendation concerns which metrics should be reported in LSM studies. As we have empirically shown here, LSM results are always spatially distorted to a certain degree – no LSM method provided output maps that were perfectly spatially accurate – so it is important to report localization using the most robust metrics. Location of Max_LSM_ or wCOM_LSM_ provided the most robust localization across LSM techniques in the current study. As also evidenced by comparison of dice scores and the OSK distribution statistic, taking statistical value intensity into account enhances the accuracy of mapping. Accordingly, using COM_LSM_ to localize the cluster or alternatively mapping localization of the whole cluster (as is often done in LSM studies) is discouraged, as these metrics are more influenced by stereotypical lesion patterns and result in mislocalization. Of all the metrics, wCOM_LSM_ is least affected by noise, while Max_LSM_ is typically closest to the underlying anatomical target. These metrics should be used when matching spatial position of LSM map clusters onto standard atlases and anatomically interpreting results.

### 5.5 Conclusions

We hope that the current systematic approach to evaluating power, accuracy and robustness of LSM methods across a range of metrics sets a new standard for these types of comparative studies. Using it as a template for future simulation analyses and validation of novel methods will help further development, understanding and proper use of LSM.

Our current results demonstrate that given sufficient lesion coverage and statistical power, both ULSM and MLSM methods are able to reliably identify distinct neural areas associated with a particular behavior or symptom. Importantly, the stereotypical lesion patterns characteristic of stroke do not prevent identification of different neural substrates for different functions under examination. However, the current study clearly showed that no LSM method, whether univariate or multivariate, was able to perfectly delineate brain regions associated with the target behavior. Simulations revealed no clear superiority of either ULSM or MLSM methods overall, but rather highlighted specific advantages of different methods. Depending on a particular study’s sample size and research questions, different types of LSM methods are more or less appropriate. Given a sufficiently large sample size, we recommend implementing both ULSM and MLSM methods with proper corrections and interpretive metrics to enhance confidence in the results. If the same anatomical foci are identified with both types of methods on the same dataset, then the results can be considered to be robust and reliable.

## Acknowledgements

We are grateful to Brian Curran and Dr. And Turken for initial assistance with collection of neuroimaging data used in the current project. We value Dr. Robert Knight’s contribution to the anatomical accuracy of lesion reconstructions over the years. We thank Dr. Stephen Wilson for developing the original VLSM software. We also extend our thanks to the developers of LESYMAP package (Dr. Dorian Pustina and others) for making their lesion database publicly available. As always, we are in debt to all the individuals with aphasia who have participated in multiple studies over the years and make this kind of work possible.

## Appendix A

**Table A1.** LSM output map location measures across different levels of behavioral noise and sample size.

|  | 0 | 0.38 | 0.77 | 32 | 48 | 64 | 80 | 96 | 112 | 128 |
| --- | --- | --- | --- | --- | --- | --- | --- | --- | --- | --- |
| **COM<sub>LSM</sub>** | 11.4 | 11.0 | 10.2 | 10.2 | 10.2 | 10.4 | 10.9 | 11.2 | 11.4 | 11.8 |
| **Max<sub>LSM</sub>** | 6.2 | 6.9 | 8.2 | 10.6 | 8.6 | 7.4 | 6.7 | 6.0 | 5.7 | 5.2 |
| **wCOM<sub>LSM</sub>** | 9.2 | 9.3 | 9.2 | 9.7 | 9.3 | 9.1 | 9.2 | 9.2 | 9.0 | 9.1 |

Notes. COM<sub>target</sub> – mean centroid location (center of mass) of the target ROI; AnyHit - nearest location of the target ROI to the LSM output map; COM<sub>LSM</sub> – mean centroid location of the LSM output; Max – maximum statistic location; wCOM<sub>LSM</sub> – mean centroid location of the LSM output weighted by statistical values.

**Table A2.** Distance-based accuracy measures with and without mask smoothing.

|  | LSM Cluster Location |  |  | Target Location |  |
| --- | --- | --- | --- | --- | --- |
|  | COM<sub>LSM</sub> | Max<sub>LSM</sub> | wCOM<sub>LSM</sub> | COM<sub>target</sub> | AnyHit |
| **0 mm** | 11 | 7.3 | 9.3 | 11.8 | 6.6 |
| **4 mm** | 10.9 | 6.8 | 9.2 | 11.5 | 6.4 |

**Table A3.**
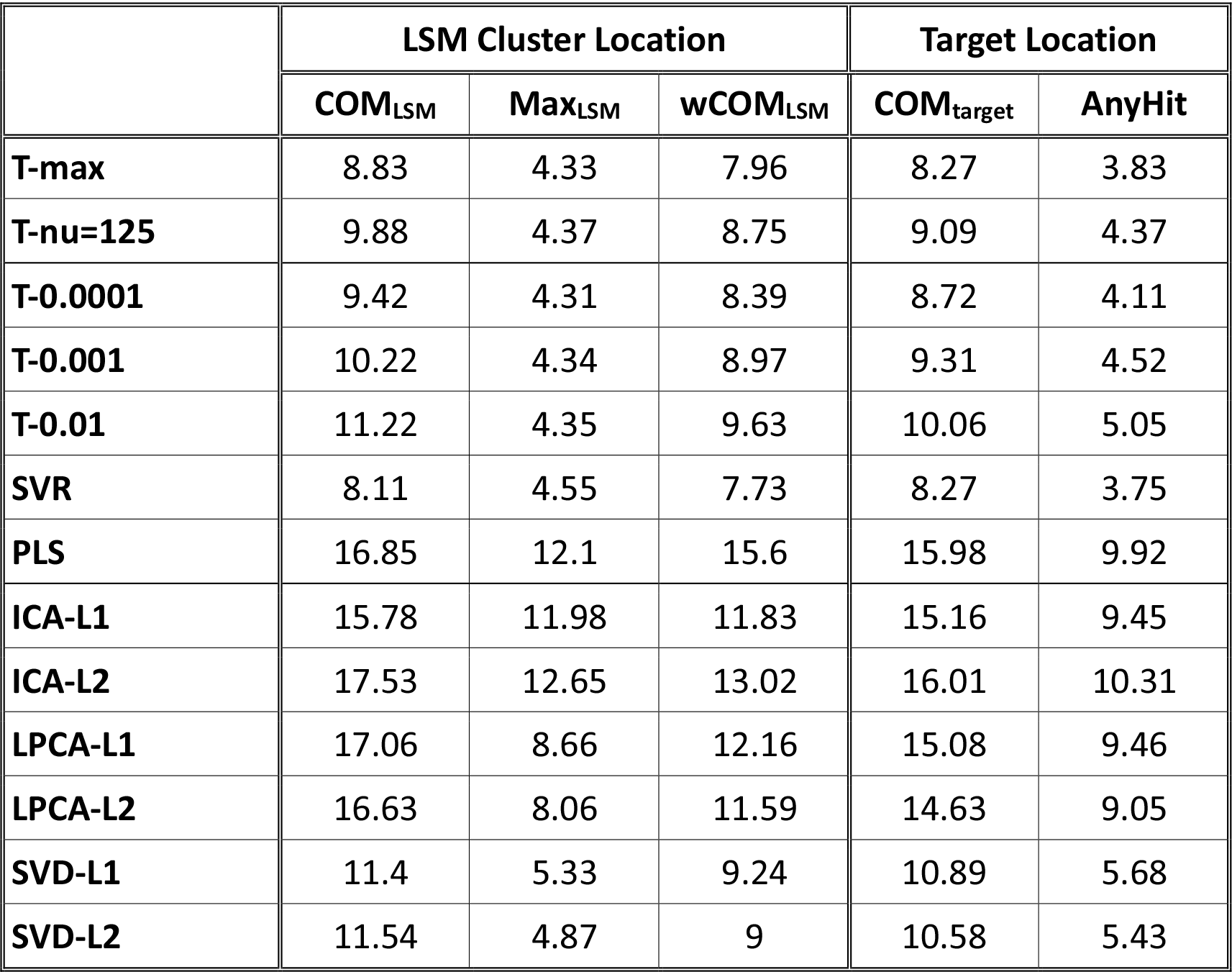
Distance-based accuracy measures across different LSM methods.

|  | LSM Cluster Location |  |  | Target Location |  |
| --- | --- | --- | --- | --- | --- |
|  | COM<sub>LSM</sub> | Max<sub>LSM</sub> | wCOM<sub>LSM</sub> | COM<sub>target</sub> | AnyHit |
| **T-max** | 8.83 | 4.33 | 7.96 | 8.27 | 3.83 |
| **T-nu=125** | 9.88 | 4.37 | 8.75 | 9.09 | 4.37 |
| **T-0.0001** | 9.42 | 4.31 | 8.39 | 8.72 | 4.11 |
| **T-0.001** | 10.22 | 4.34 | 8.97 | 9.31 | 4.52 |
| **T-0.01** | 11.22 | 4.35 | 9.63 | 10.06 | 5.05 |
| **SVR** | 8.11 | 4.55 | 7.73 | 8.27 | 3.75 |
| **PLS** | 16.85 | 12.1 | 15.6 | 15.98 | 9.92 |
| **ICA-L1** | 15.78 | 11.98 | 11.83 | 15.16 | 9.45 |
| **ICA-L2** | 17.53 | 12.65 | 13.02 | 16.01 | 10.31 |
| **LPCA-L1** | 17.06 | 8.66 | 12.16 | 15.08 | 9.46 |
| **LPCA-L2** | 16.63 | 8.06 | 11.59 | 14.63 | 9.05 |
| **SVD-L1** | 11.4 | 5.33 | 9.24 | 10.89 | 5.68 |
| **SVD-L2** | 11.54 | 4.87 | 9 | 10.58 | 5.43 |

**Table A4.** Average displacement error (mm) across all LSM methods when using and not using lesion size as a covariate.

|  | Lesion size covariate | No lesion size covariate |
| --- | --- | --- |
| **T-max** | 5.2 | 7.3 |
| **T-nu=125** | 6.1 | 8.4 |
| **T-0.0001** | 5.7 | 7.9 |
| **T-0.001** | 6.3 | 8.9 |
| **T-0.01** | 7.2 | 10.5 |
| **SVR** | 5.3 | 8.0 |
| **PLS** | 12.0 | 20.2 |
| **ICA-L1** | 12.4 | 14.7 |
| **ICA-L2** | 13.8 | 15.7 |
| **LPCA-L1** | 12.4 | 16.1 |
| **LPCA-L2** | 11.6 | 19.2 |
| **SVD-L1** | 8.2 | 11.9 |
| **SVD-L2** | 7.2 | 10.8 |

**Table A5.**
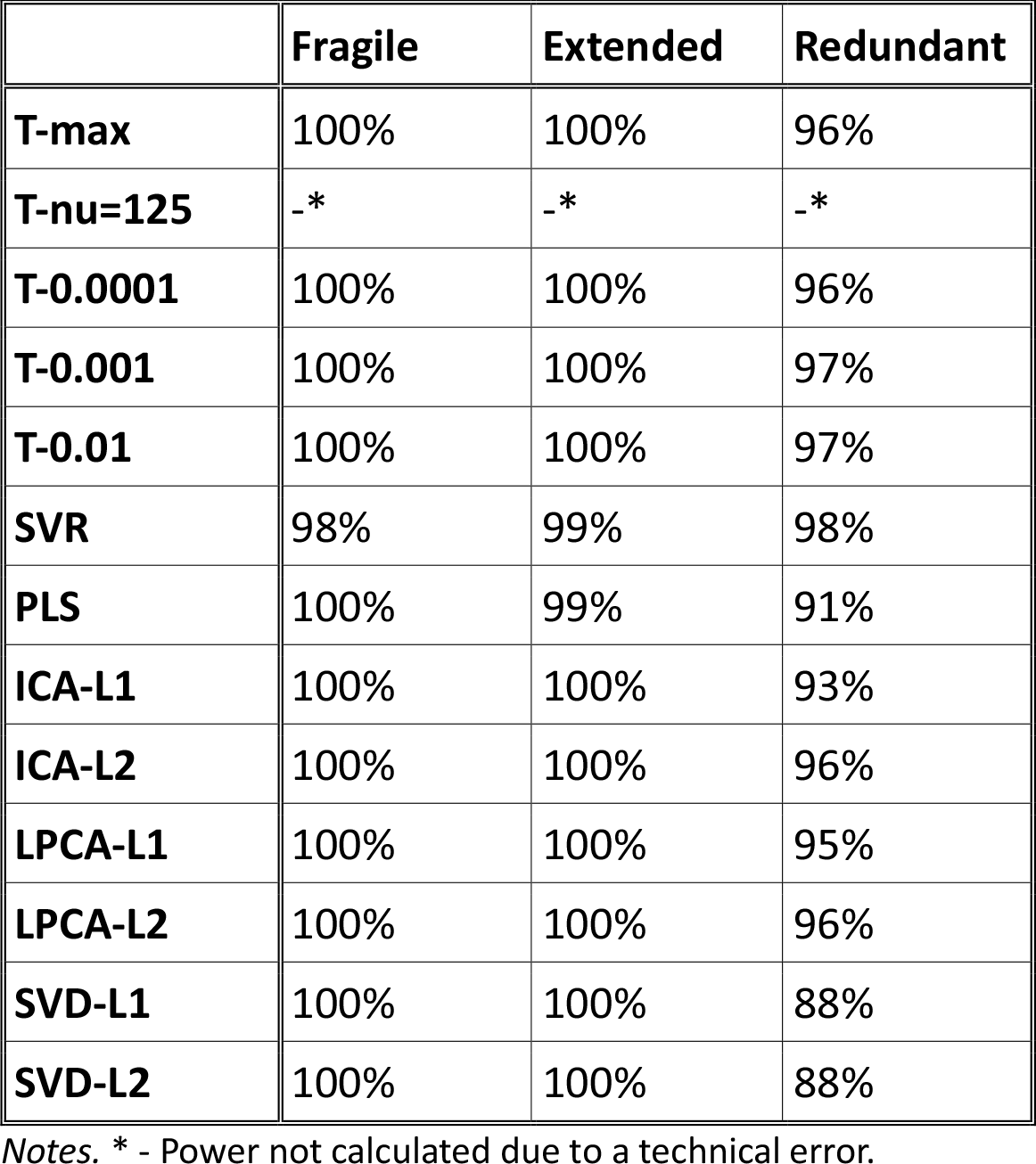
Power (percentage of trials where significant voxels were detected) for dual target simulations with three networks across LSM methods.

|  | Fragile | Extended | Redundant |
| --- | --- | --- | --- |
| **T-max** | 100% | 100% | 96% |
| **T-nu=125** | _* | _* | _* |
| **T-0.0001** | 100% | 100% | 96% |
| **T-0.001** | 100% | 100% | 97% |
| **T-0.01** | 100% | 100% | 97% |
| **SVR** | 98% | 99% | 98% |
| **PLS** | 100% | 99% | 91% |
| **ICA-L1** | 100% | 100% | 93% |
| **ICA-L2** | 100% | 100% | 96% |
| **LPCA-L1** | 100% | 100% | 95% |
| **LPCA-L2** | 100% | 100% | 96% |
| **SVD-L1** | 100% | 100% | 88% |
| **SVD-L2** | 100% | 100% | 88% |

Notes. \* - Power not calculated due to a technical error.

**Table A6.** Power (percentage of trials where significant voxels were detected) for dual target simulations with three networks across sample sizes.

|  | **Fragile** | **Extended** | **Redundant** |
| --- | --- | --- | --- |
| **64** | 98% | 98% | 82% |
| **80** | 99% | 100% | 88% |
| **96** | 100% | 100% | 93% |
| **112** | 100% | 100% | 95% |
| **128** | 100% | 100% | 97% |
| **144** | 100% | 100% | 97% |
| **160** | 100% | 100% | 98% |
| **176** | 100% | 100% | 99% |
| **192** | 100% | 100% | 99% |
| **208** | 100% | 100% | 99% |

**Table A7.** Dice coefficient values for dual target simulations with three networks across LSM methods.

|  | **Fragile** | **Extended** | **Redundant** |
| --- | --- | --- | --- |
| **T-max** | 0.13 | 0.13 | 0.13 |
| **T-nu=125** | 0.11 | 0.11 | 0.11 |
| **T-0.0001** | 0.12 | 0.12 | 0.12 |
| **T-0.001** | 0.1 | 0.11 | 0.11 |
| **T-0.01** | 0.08 | 0.08 | 0.09 |
| **SVR** | 0.14 | 0.15 | 0.13 |
| **PLS** | 0.05 | 0.05 | 0.05 |
| **ICA-L1** | 0.07 | 0.07 | 0.07 |
| **ICA-L2** | 0.06 | 0.06 | 0.06 |
| **LPCA-L1** | 0.06 | 0.07 | 0.06 |
| **LPCA-L2** | 0.06 | 0.07 | 0.07 |
| **SVD-L1** | 0.07 | 0.06 | 0.07 |
| **SVD-L2** | 0.07 | 0.07 | 0.08 |

